# Modeling the effect of cooperativity in ternary complex formation and targeted protein degradation mediated by heterobifunctional degraders

**DOI:** 10.1101/2022.03.22.485399

**Authors:** Daniel Park, Jesus Izaguirre, Rory Coffey, Huafeng Xu

**Affiliations:** Roivant Discovery, New York, New York 10036, United States

## Abstract

Chemically induced proximity between certain endogenous enzymes and a protein of interest (POI) inside cells may cause post-translational modifications to the POI with biological consequences and potential therapeutic effects. Heterobifunctional (HBF) molecules that bind with one functional part to a target POI and with the other to an E3 ligase induce the formation of a target-HBF-E3 ternary complex, which can lead to ubiquitination and proteasomal degradation of the POI. Targeted protein degra-dation (TPD) by HBFs offers a promising approach to modulating disease-associated proteins, especially those that are intractable using other therapeutic approaches, such as enzymatic inhibition. The three-way interactions among the HBF, the target POI, and the ligase—including the protein-protein interaction (PPI) between the POI and the ligase—contribute to the stability of the ternary complex, manifested as positive or negative binding cooperativity in its formation. How such cooperativity affects HBF-mediated degradation is an open question. In this work, we develop a pharmacodynamic model that describes the kinetics of the key reactions in the TPD process, and we use this model to investigate the role of cooperativity in the ternary complex formation and in the target POI degradation. Our model predicts that, under certain conditions, increasing cooperativity may diminish degradation, implying an optimal range of cooperativity values for efficient degradation. We also develop a statistical inference model for determining cooperativity in intracellular ternary complex formation from cellular assay data, and demonstrate it by quantifying the change in cooperativity due to site-directed mutagenesis at the POI-ligase interface of the SMARCA2-ACBI1-VHL ternary complex. Our pharmacodynamic model provides a quantitative framework to dissect the complex HBF-mediated TPD process and may inform the rational design of effective HBF degraders.

## Introduction

Targeted protein degradation (TPD) mediated by heterobifunctional (HBF) degraders has emerged as a promising modality for therapeutic applications,^1–3^ which enables in-cell reduction of a target POI that might not be amenable to traditional stoichiometric inhibition. HBF degraders, such as proteolysis targeting chimeras (PROTACs),^4^ simultaneously bind to an E3 ligase and a target POI, resulting in a target-HBF-E3 ternary complex,^5–7^ in which the E3 ligase recruits a ubiquitin-charged E2 enzyme to ubiquitinate the POI, labeling the POI with a polyubiquitin chain for subsequent degradation by the proteasome.^8^

The formation of the ternary complex often exhibits cooperativity in that the free energy change attendant to its formation is more negative or more positive than the sum of the free energy changes associated with the formation of the binary complexes between the E3 ligase and the HBF and between the target POI and the HBF.^9,10^ The cooperativity—denoted as *α*—is defined to be the ratio of the equilibrium association constant of the E3 ligase binding to the HBF-target binary complex to that of the E3 ligase binding to the HBF alone;^10^ by thermodynamic symmetry, it is the same as the ratio of the equilibrium constant of the target binding to the HBF-E3 binary complex to that of the target binding to the HBF alone. Positive cooperativity (*α >* 1) increases and negative cooperativity (*α <* 1) decreases the thermodynamic stability of the ternary complex. Among other reasons for cooperativity, the E3 ligase and the target POI form a novel protein-protein interface in the ternary complex, which may be thermodynamically favorable or unfavorable. Different HBF molecules may beget different cooperativity for the same pair of target POI and E3 ligase, ^7^ providing a route to optimize the HBF molecule for efficient ternary complex formation and target degradation.^6^ Conversely, an HBF may induce different cooperativity between two very similar POIs when forming a ternary complex with the same E3 ligase, ^5,11^ giving rise to the opportunity of designing selective degraders using a promiscuous warhead molecule. ^12,13^ Because of the critical role of the ternary complex in the TPD process, cooperativity is an important parameter to consider in the design of HBF degraders.

Although experimental evidence so far suggests that stable ternary complexes typically lead to efficient degradation,^14–17^ there have been conflicting reports on the role of cooperativity in HBF-mediated TPD.^9,18^ Strong degraders have been reported to have a wide range of cooperativity values,^10^ including negative cooperativity.^19,20^ Conversely, HBF molecules with high cooperativity may be ineffective degraders.^7^ Unraveling the role of cooperativity in degradation requires a mechanistic model that relates the stability of the ternary complex to degradation efficiency.

In this work, we develop a quantitative model of HBF-mediated TPD that predicts degradation efficacy given the kinetic parameters of the key steps in the TPD process, including those associated with ternary complex formation. Using this model, we investigate the effect of varying cooperativity—which varies the ternary complex stability while holding the other factors constant—on the degradation efficiency. Our analysis suggests that there may exist conditions under which increasing cooperativity (hence the ternary complex stability) diminishes degradation, and the optimal value of cooperativity to maximize degradation depends on other parameters, such as the stability of the binary complexes, the ubiquitination rate, the degradation rate, the expression level of the target POI, and the time point of degradation measurements. The predictions by our model on how degradation changes in response to changing parameters can be tested experimentally.

In addition, we develop a statistical inference model using Markov chain Monte Carlo (MCMC) to estimate the cooperativity value from in-cell ternary complex formation assays. As an example, we have used it and the NanoBRET assay^21^—which measures the relative intracellular ternary complex concentrations at different HBF concentrations—to determine the cooperativity of the ternary complex formed by the SMARCA2 protein, the HBF molecule ACBI1,^6^ and the von Hippel-Lindau (VHL) E3 ligase, including SMARCA2 and VHL mutants engineered by site-directed mutagenesis to potentially perturb their PPI in the ternary complex. Our analysis quantifies the contribution of different specific interactions at the protein-protein interface to the stability of the ternary complex inside the cell.^6^

The complexity of the multi-step TPD process challenges the empirical approach to degrader development. The degradation efficiency depends on many parameters and, as we demonstrate below, the effect of modulating one parameter depends on the values of the other parameters. It is often unclear why an HBF molecule induces good or poor target degradation. By identifying the key parameters to which degradation is most sensitive under different biological contexts and predicting how degradation responds to changes in the parameter values, our model may help guide the development of effective HBF degraders.

## Methods

### Theoretical Models

#### Pharmacodynamic model of HBF-mediated protein degradation

We have devised a pharmacodynamic model of HBF-mediated TPD using a system of time-dependent ordinary differential equations to describe the kinetics of the reactions involved in the degradation process. The model simulates the following reactions that occur once a dose of HBF molecules is introduced into the extracellular environment (Tables 1 and 2, Fig. 1): The HBF molecule traverses the cell membrane (Reaction 1); HBF forms binary complexes with either the target POI (Reaction 2) or the E3 ligase (Reaction 3); A ternary complex forms (via Reactions 4 or 5); The E3 ligase in the ternary complex recruits ubiquitin-charged E2 enzymes to ubiquitinate the target (Reaction 6); The ubiquitinated target dissociates from the ternary complex with HBF bound (Reverse reactions 4) or without (Reverse reaction 5); The ubiquitinated target (or the HBF-target binary complex) either undergoes a deubiquitination cascade (Reaction 7), or if the target is polyubiquitinated with a chain of *n* = 4 ubiquitin molecules, it is degraded by the proteasome (Reaction 8); Meanwhile, the target POI is produced by protein synthesis (Reaction 10) and targets in all forms undergo intrinsic, HBF-independent degradation (Reaction 11). In the initial HBF-free state, the baseline target concentration is determined by the ratio of the target production rate to the intrinsic degradation rate, satisfying

**Table 1:** The molecular species in our model of TPD. In the following, we will use *T*_*i*_ · * to denote the sum of *T*_*i*_ in free, binary, and ternary forms, and 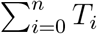 to denote the sum of free target POI in all ubiquitination states.

| Notation | Species |
| --- | --- |
| $D_{ec}$ | Extracellular HBF degrader |
| $D_{ic}$ | Intracellular HBF degrader |
| $T_i$ | Free target with ubiquitin chain of length $i$; $T_0$ denotes unmodified target |
| $E3$ | Free E3 ligase |
| $T_i \cdot D$ | Binary complex of target and HBF |
| $D \cdot E3$ | Binary complex of HBF and E3 |
| $T_i \cdot D \cdot E3$ | Ternary complex of target, HBF, and E3 |

**Table 2:** Reactions in our model of HBF-mediated target protein degradation.

|  |  |  |
| --- | --- | --- |
| 1 | $D_{ec} \xrightleftharpoons[PS_{cell}]{PS_{cell}} D_{ic}$ | HBF permeation and efflux in/out of the cell |
| 2 | $D_{ic} + T_i \xrightleftharpoons[k_{off,T,binary}]{k_{on,T,binary}} T_i \cdot D$ | HBF-target binary complex formation |
| 3 | $D_{ic} + E3 \xrightleftharpoons[k_{off,E3,binary}]{k_{on,E3,binary}} D \cdot E3$ | HBF-E3 binary complex formation |
| 4 | $T_i \cdot D + E3 \xrightleftharpoons[k_{off,E3,ternary}]{k_{on,E3,ternary}} T_i \cdot D \cdot E3$ | Ternary complex formation |
| 5 | $T_i + D \cdot E3 \xrightleftharpoons[k_{off,T,ternary}]{k_{on,T,ternary}} T_i \cdot D \cdot E3$ | Ternary complex formation |
| 6 | $T_i \cdot D \cdot E3 \xrightarrow{k_{ub}} T_{i+1} \cdot D \cdot E3$ | Target ubiquitination in ternary complex |
| 7 | $T_i \text{ (or } T_i \cdot D) \xrightarrow{k_{deub}} T_{i-1} \text{ (or } T_{i-1} \cdot D)$ | Target deubiquitination |
| 8 | $T_n \text{ (or } T_n \cdot D) \xrightarrow{k_{deg,T}} \emptyset \text{ (or } D_{ic})$ | Polyubiquitinated target degradation |
| 9 | $T_n \cdot D \cdot E3 \xrightarrow{k_{deg,Ternary}} D \cdot E3$ | Polyubiquitinated ternary complex degradation |
| 10 | $\emptyset \xrightarrow{k_{prod}} T_0$ | Target synthesis |
| 11 | $T_i \cdot * \xrightarrow{k_{deg,int}} \emptyset \text{ for } i = 0, 1, \dots, n$ | Intrinsic target degradation |

**Figure 1:**
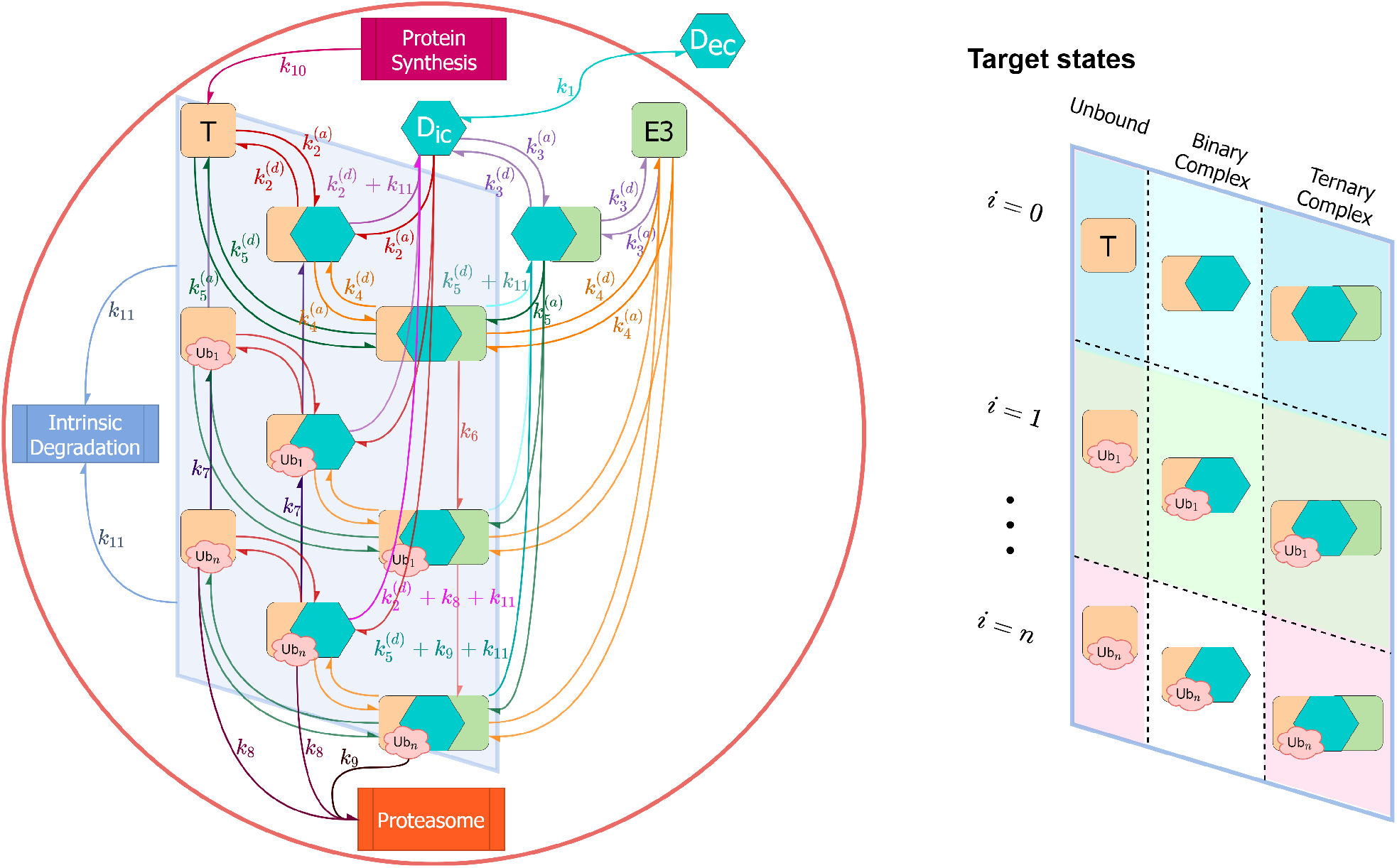
Schematic of our TPD model. The molecular species in our model are summarized in Table 1. Rate constants, defined in Tables 3 and 4, are abbreviated as follows: *k*_1_ = Permeability surface area product, 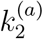 = Target binary association rate, 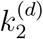 = Target binary dissociation rate, 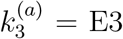 binary dissociation rate, 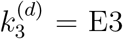 binary dissociation rate 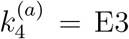 ternary association rate, 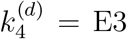 ternary dissociation rate, 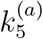 = Target ternary association rate, 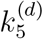 = Target ternary dissociation rate, *k*_6_ = Target ubiquitination rate, *k*_7_ = Target deubiquitination rate, *k*_8_ = Polyubiquitinated target proteasomal degradation rate, *k*_9_ = Polyubiquitinated ternary complex proteasomal degradation rate, *k*_10_ = Target synthesis rate, *k*_11_ = Intrinsic target degradation rate.

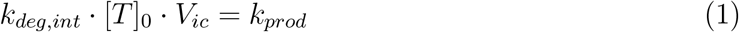

**Table 3:** Reference parameters for TPD simulations.

| Parameter | Value | Notes |
| --- | --- | --- |
| *Ubiquitination and Degradation* |  |  |
| $n$ | 4 | Length of polyubiquitin chain on POI required for degradation |
| $k_{ub}$ | 1500/h | $k_6$ in Figure 1; Previously reported simulation values for $k_{ub}$ have ranged from 6/h<sup>23</sup> to 2667/h<sup>22</sup> |
| $k_{deub}$ | 360/h | $k_7$ in Figure 1 |
| $k_{deg,T}$ | 60/h | $k_8$ in Figure 1 |
| $k_{deg,Ternary}$ | 60/h or 0/h | $k_9$ in Figure 1 |
| *Physiological System Parameters* |  |  |
| $f_{u_{ec}}$ | 1 | Extracellular HBF degrader fraction unbound |
| $f_{u_{ic}}$ | 1 | Intracellular HBF degrader fraction unbound |
| $k_{prod}$ | $5.24 \times 10^{-15}$ $\mu\text{mol/h}$ | $k_{10}$ in Figure 1; Calculated by Eq. 1 |
| $k_{deg,int}$ | 0.1/h | $k_{11}$ in Figure 1 |
| $[T]_0$ | 1 $\mu\text{M}$ | Baseline target protein concentration |
| $[E3]_0$ | 1 $\mu\text{M}$ | Baseline E3 ligase concentration |
| $[D]_0$ | 1 nM | Baseline extracellular degrader exposure |
| $num_{cells}$ | 5000 | Number of cells |
| $V_{ic}$ | $5.24 \times 10^{-13}$ L | Intracellular volume assuming a spherical cell with 10 $\mu\text{m}$ diameter |
| $V_{ec}$ | $2 \times 10^{-4}$ L | Extracellular volume |

**Table 4:** Molecule-specific ternary complex formation and permeability reference parameters for TPD simulations.

| Parameter | ACBI1 | PROTAC 1 | Notes |
| --- | --- | --- | --- |
| $\alpha$ | 26 | 3.2 | |
| $k_{on,T,binary}$ | 36000/μM/h | 36000/μM/h | $k_2^{(a)}$ in Figure 1 |
| $k_{off,T,binary}$ | 333360/h | 307000/h | $k_2^{(d)}$ in Figure 1 |
| $K_{D,T,binary}$ | 9.26 μM | 8.54 μM | |
| $k_{on,E3,binary}$ | 5364/μM/h | 8929/μM/h | $k_3^{(a)}$ in Figure 1 |
| $k_{off,E3,binary}$ | 372.26/h | 109.8/h | $k_3^{(d)}$ in Figure 1 |
| $K_{D,E3,binary}$ | 0.07 μM | 0.01 μM | |
| $k_{on,T,ternary}$ | 878.4/μM/h | 1026/μM/h | $k_5^{(a)}$ in Figure 1 |
| $k_{off,T,ternary}$ | 312.85/h | 2721.6/h | $k_5^{(d)}$ in Figure 1 |
| $K_{D,T,ternary}$ | 0.36 μM | 2.66 μM | |
| $k_{on,E3,ternary}$ | 878.4/μM/h | 1026/μM/h | $k_4^{(a)}$ in Figure 1 |
| $k_{off,E3,ternary}$ | 2.34/h | 3.94/h | $k_4^{(d)}$ in Figure 1 |
| $K_{D,E3,ternary}$ | 0.0027 μM | 0.0038 μM | |
| $PS_{cell}$ | $2.49 \times 10^{-11}$ L/h | $1.24 \times 10^{-12}$ L/h | $k_1$ in Figure 1 Permeability surface area using reported apparent permeability<sup>6</sup> multiplied by surface area assuming spherical cell with 10 μm diameter |

The reference simulation parameters used in this work are listed in Table 3, and exemplary parameters of complex formation and membrane permeability for two HBF molecules are listed in Table 4. Mass action kinetics^22^ govern the interactions between the target POI, HBF, and E3 ligase during ternary complex formation, ubiquitination, deubiquitination, and degradation. We assume that the baseline total E3 ligase concentration stays constant.

Our model is derived from previous models of HBF-mediated protein degradation,^22,23^ but with important modifications, which enable us to reach different qualitative conclusions than previous models. First, our model accounts for a deubiquitination cascade after the ubiquitinated target protein dissociates from the ternary complex, whereas previous models assume instantaneous detachment of the entire ubiquitin chain from the target protein upon dissociation. Explicit consideration of the deubiquitination cascade results in an additional set of molecular species in our model that are absent in the previous models: the free target protein in various stages of polyubiquitination, which may be either rescued from degradation by the finite-rate deubiquination process, re-engaged in the ternary complex with its ubiquitin chain further extended, or, for the protein with the ubiquitin chain length of *n* = 4, degraded. Second, our model assumes that the polyubiquitinated target protein may be degraded by the proteasome at a different—presumably no slower—rate after it dissociates from the ternary complex than when it is in a ternary complex (*k*_*deg,T*_ *> k*_*deg,T ernary*_), instead of always setting the same degradation rate for the polyubiquitinated target regardless whether it is free or bound in the ternary complex.^22^ This assumption is only applicable in our model because the previous models, as mentioned above, do not admit the free polyubiquitinated target protein. In this work, we first consider two limiting scenarios: 1) *k*_*deg,T ernary*_ = *k*_*deg,T*_, in which the proteasome degrades the target in the ternary complex as efficiently as the free target, and 2) *k*_*deg,T ernary*_ = 0, in which the proteasome does not degrade the target in the ternary complex (Table 3; see Discussion). We will then explore how the relationship between degradation and cooperativity is affected by the ratio of *k*_*deg,T ernary*_*/k*_*deg,T*_.

#### Determining cooperativity from intracellular assay of ternary complex formation

The equilibrium ternary complex concentration can be calculated by solving the equilibrium mass action and conservation of mass equations,^24,25^ if the binary binding constants, cooperativity, and the concentrations of HBF, target POI, and E3 ligase are known. Common, however, is the inverse problem: determining the values of these parameters from the experimental measurements of ternary complex formation at different HBF concentrations. In experiments that measure *intracellular* ternary complex concentrations,^21^ the intracellular concentrations of HBF, target POI, and E3 ligase are *a priori* unknown, and the cooperativity may differ from that measured in a biophysical assay because the intracellular target protein is in a different environment and may be in a different state of conformation or post-translational modification, which make it challenging to robustly determine all these parameters at once. Here, we develop a model within the Bayesian framework of statistical inference to estimate these parameters via MCMC in fitting the equilibrium concentration equations to the experimentally measured intracellular ternary complex formation.

MCMC is an iterative procedure that samples from a posterior probability distribution of parameters given the observed data and prior probabilistic assumptions about the parameters.^26,27^ While exploring the target parameter space through sampling, Bayesian inference is used to update the prior state of beliefs such that the likelihood of the observed data is maximized by the resulting posterior state of beliefs.^28^ In cases when the posterior probability distribution is analytically intractable, MCMC provides repeated random samples from the posterior, from which characteristics of the distribution such as the mean and highest density intervals can be approximated.

A schematic of our probabilistic graphical model is shown in Fig. 2, and the model parameters are listed in Table 5. Experiments such as the NanoBRET assay^21^ measure the intracellular ternary complex formation by a readout that is proportional to the ternary complex concentration. Given the readouts measured at *N* different extracellular HBF concentrations [*D*_*ec*_], our model estimates the posterior probability distributions of the unknown parameters in the ternary complex equilibrium. In this work, we apply our model to simultaneously determine the parameters of *M* = 5 ternary complexes, which share the same HBF molecule but with either the target protein or the E3 ligase mutated at the protein-protein interface, presumably altering the cooperativity. In our analysis, we assume that all constructs share the same binary binding constants but have different cooperativity values {*α*_*i*=1,2,…,*M*_}, and that at equilibrium the intracellular HBF concentration is proportional to the initial extracellular concentration: [*D*_*ic*_]_*t*_ = *κ*[*D*_*ec*_]. The readout (milliBRET unit, or mBU, in NanoBRET assay) is assumed to be proportional to the ternary complex concentration: *mBU* = exp(*β*)[*T* · *D* · *E*3], where exp(*β*) is the proportionality constant. The experimentally observed readout is taken to be normally distributed, centered at the predicted *mBU* with standard deviation *σ*. At each iteration, MCMC proposes candidate values for the total target concentration [*T*]_*t*_, the total E3 ligase concentration [*E*3]_*t*_, cooperativity {*α*_*i*_}, *κ, β*, and *σ*. Given these proposed values and fixed binary equilibrium dissociation constants, the equilibrium ternary complex concentration [*T* · *D* · *E*3] is computed by numerically solving the mass action and mass conservation equations. The proposed values are then accepted or rejected according to the Metropolis criteria based on the likelihood of the observed readouts.^27^ We ran 4 independent MCMC chains at different initial parameter values, each for 10,000 iterations. The first 1,000 samples were discarded (burn-in), as was every second sample thereafter (thinning). The {*α*_*i*_} parameters showed good convergence, achieving Gelman-Rubin convergence diagnostic statistics^29^ of 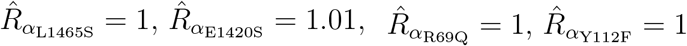.

**Table 5:** Parameters in ternary complex equilibrium model.

| Parameter | Prior | Description |
| --- | --- | --- |
| *Unknown Variables to Estimate* |  |  |
| $\kappa$ | $\mathcal{U}[0,1]$ | Permeability proportion at equilibrium |
| $[T]_t$ | $\text{Exp}(1)$ | Total free target concentration. Posterior mean: 13 nM.<br>95% credible interval: [4 nM, 33 nM]. |
| $[E3]_t$ | Exp(1) | Total E3 ligase concentration. Posterior mean: 34 nM. 95% credible interval: [12 nM, 84 nM]. |
| $\alpha_i$ | Lognormal(3, $0.5^2$) | Cooperativity of ternary complex type $i$; $\alpha$ for wild type SMARCA2, VHL, and ACBI1 fixed at 26, within previously reported values<sup>6</sup> |
| $\beta$ | $\mathcal{N}(0, 10^2)$ | Ternary complex concentration to mBU scaling factor |
| $\sigma$ | Exp(10) | Standard deviation of likelihood |
| *Data and Fixed Quantities* |  |  |
| $[D_{ec}]$ | | Total extracellular degrader concentration |
| $K_{D,T}$ | | Equilibrium dissociation constant of HBF-target binary complex; $K_{D,T} = 1.8 \mu\text{M}$ for ACBI1-SMARCA2<sup>6</sup> |
| $K_{D,E3}$ | | Equilibrium dissociation constant of HBF-E3 binary complex; $K_{D,E3} = 0.25 \mu\text{M}$ for ACBI1-HBF<sup>6</sup> |
| *Deterministic Quantities* |  |  |
| $[D_{ic}]_t$ | | Total intracellular degrader concentration |
| $[T]$ | | Free target concentration at equilibrium |
| $[E3]$ | | Free E3 ligase concentration at equilibrium |
| $[T \cdot D \cdot E3]$ | | Ternary complex concentration at equilibrium |
| *Likelihood* |  |  |
| mBU | | Assay response; $\mathcal{N}([T \cdot D \cdot E3], \sigma^2)$ |

**Figure 2:**
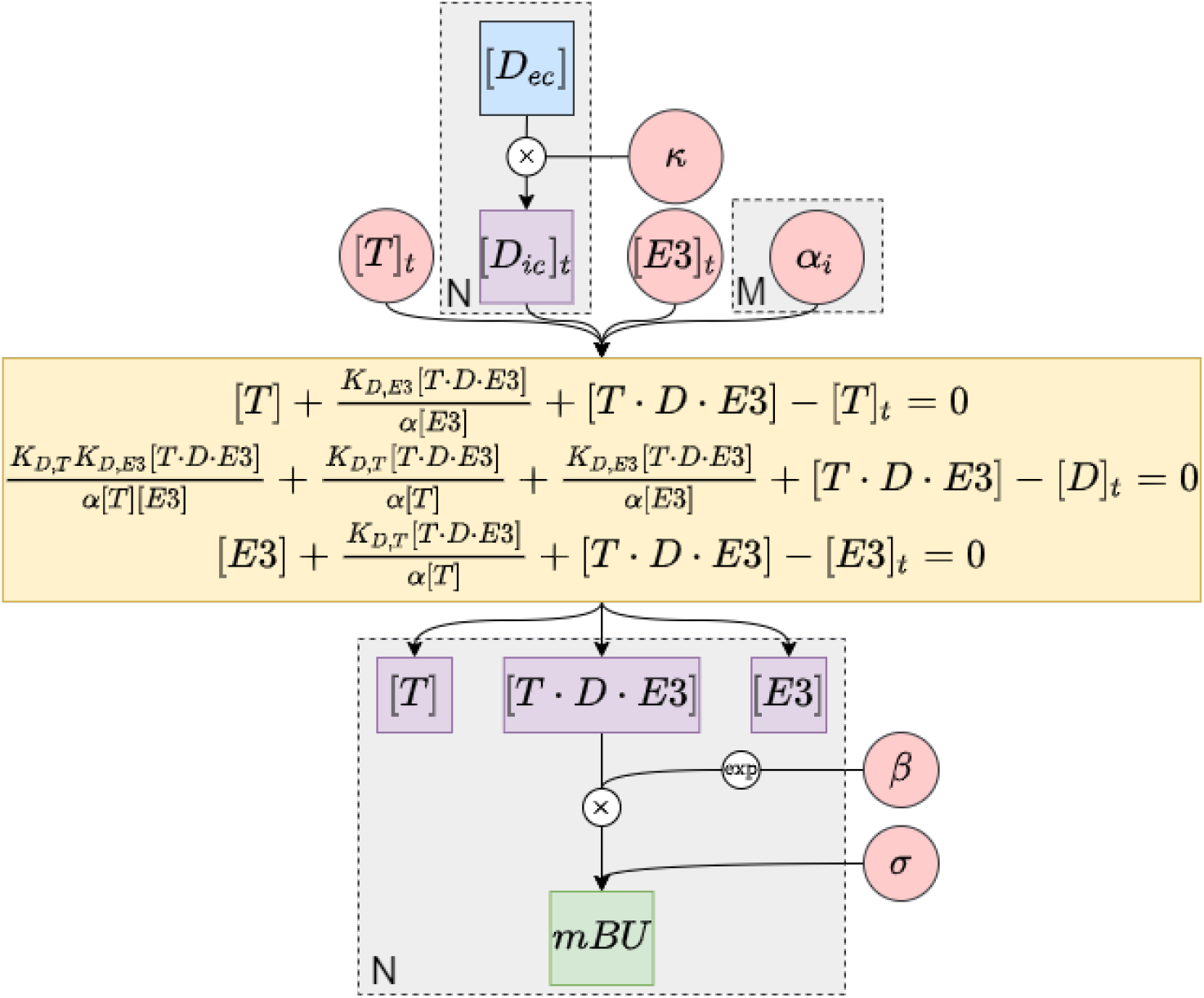
Bayesian belief network showing the conditional dependencies between the parameters in a ternary complex equilibrium system. Blue nodes = observed data, Red nodes = parameters to estimate, Purple nodes = deterministic quantities, Green nodes = likelihood of observed data. Plate notation denotes repeated nodes: *N* = total number of ternary complex formation observations, *M* = number of constructs. Model parameters are defined in Table 5. System of equations (yellow) describing molecular species concentrations at equilibrium is taken from Douglass *et al*.^24^ and solved numerically.

## Experimental Procedures

### Plasmids

Mammalian expression vectors

- pRP-CMV-HaloVHL-IRES-nLucSMARCA2
- pRP-CMV-HaloVHL(R69Q)-IRES-nLucSMARCA2
- pRP-CMV-HaloVHL(Y112F)-IRES-nLucSMARCA2
- pRP-CMV-HaloVHL-IRES-nLucSMARCA2(E1420S)
- pRP-CMV-HaloVHL-IRES-nLucSMARCA2(L1465S)
- pRP-EF1A-EloB-IRES-EloC

were ordered and cloned by VectorBuilder (https://en.vectorbuilder.com/). Sequences of the genes are listed in Supplementary Information.

### NanoBRET assay

HEK293T (ATCC CRL-3216) cells were maintained in 37 °C incubators with 5% CO_2_ and grown in complete media, DMEM (Corning 10-013-CM) supplemented with 10% FBS (Corning 35-011-CV), and 1% L-glutamine (Corning glutagro 25-015-CL). For transfections, HEK293T cells were seeded at 5×10^5^ per well in a 6-well plate (day 1). 24 hours post seeding (day 2), lipofectamine 2000 (ThermoFisher Scientific 11668019) was used as per manufacturer’s protocol to co-transfect 2 *µ*g of the pRP-CMV-HaloVHL-IRES-nLucSMARCA2 and 2 *µ*g of the pRP-EF1A-EloB-IRES-EloC plasmid. 24 hours after transfection (day 3), cells were removed from the 6-well plates by trypsinization (0.25% Trypsin EDTA Gibco 25200-056) and seeded into a 96 well cell culture treated flat bottom white plate (Corning 3917) in 100 *µ*L per well with phenol red free Opti-MEM (Gibco 11058-021) supplemented with 4% FBS and 100 nM HaloTag NanoBRET 618 ligand (Promega N2583). Additional cells were plated without the HaloTag NanoBRET 618 ligand to serve as background control. 24 hours after seeding into white bottom plates (day 4), half of the media was removed and replaced with compounds for dose response testing with the final makeup of the media being 0.1% DMSO, 2 *µ*M Mg132 (Cell Signaling 2194S), 1:100 Vivazine (Promega N2583), and 4% FBS in phenol red free Opti-MEM. The plates were immediately measured using a Biotek Synergy Neo2 at 450 and 610 nM with Biotek 74 NanoBRET Dual PTM/LUM upper top filter and read at various time points thereafter. We fit our model to the readings at 97 min.

### Data Availability

All data and code will be made available on GitHub upon the publication of the manuscript.

## Results

### The effect of cooperativity on degradation

To illustrate the effect of cooperativity on degradation, we compare two HBF molecules with similar binary affinities for the POI and for the E3 ligase, but with about an order of magnitude difference in their cooperativity values. One such example is the pair of ACBI1 and PROTAC 1, both HBF molecules developed to recruit the E3 ligase VHL for the degradation of the target protein SMARCA2.^6^ ACBI1 induces a more stable ternary complex than PROTAC 1 does, due to a higher cooperativity (26 for ACBI1 vs 3.2 for PROTAC 1), even though ACBI1 has a weaker affinity for both the target protein (SMARCA2) and the E3 ligase (VHL) (Table 4). ACBI1 has been reported to degrade SMARCA2 more effectively than PROTAC 1 does.^6^

Using our model, we simulated the target degradation induced by these two molecules, guess-estimating the experimentally unavailable kinetic parameters associated with ternary complex formation and with the degradation process (Table 3 and 4).^22^ Our simulations recapitulate the experimental observation that ACBI1 is a stronger degrader than PROTAC 1 (Fig. 3a). To control for the effect of permeability (ACBI1 is 20 times more permeable than PROTAC 1^6^), we simulated a variant of PROTAC 1, denoted by PROTAC 1*, with its permeability set to that of ACBI1. ACBI1 induces quicker degradation than PROTAC 1* (Fig. 3a), even though the intracellular concentration of PROTAC 1* is higher than that of ACBI1 (Fig. 3b). ACBI1 degrades the target faster than PROTAC 1* (Fig. 3a) because the former induces a higher concentration of ternary complex (due to its higher cooperativity), leading to a faster accumulation of polyubiquitinated target primed for degradation (Fig. 4). At thermodynamic equilibrium, ternary complex formation first increases with increasing degrader concentration but then decreases as the degrader concentration increases beyond a certain value (Fig. 5). This is known as the hook effect:^24^ at sufficiently high degrader concentration, different HBF molecules will form binary complexes with either the POI or the E3, yet no HBF molecule will bridge the POI and the E3 to form the ternary complex.

**Figure 3:**
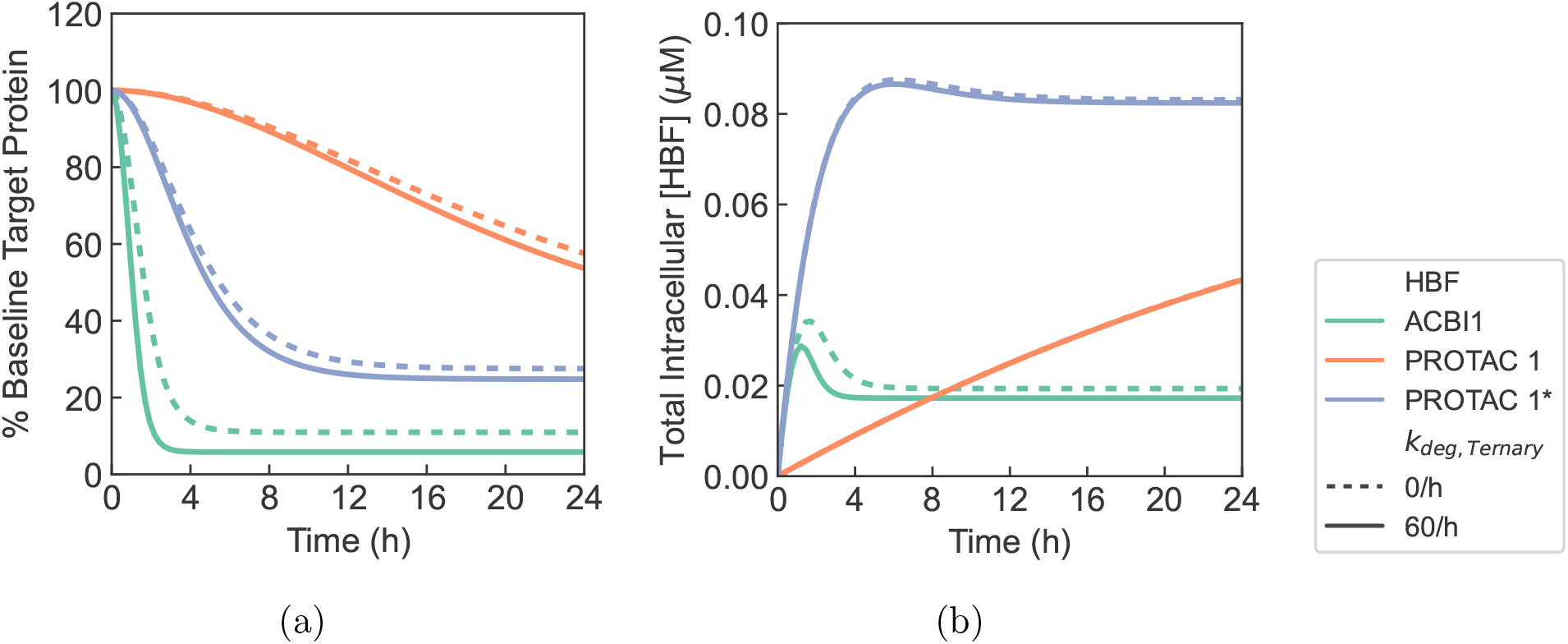
(a) Simulated degrader-induced target protein degradation of SMARCA2. (b) Total intracellular HBF concentration, [*D*_*ic*_ · *], over time. Permeability of PROTAC 1 was set to the *PS*_*cell*_ parameter value for PROTAC 1 in Table 4, and permeability of PROTAC 1* was set to the *PS*_*cell*_ value for ACBI1 in Table 4. Two degradation rates of the ternary complex are considered: *k*_*deg,T ernary*_ = *k*_*deg,T*_ = 60*/*h and *k*_*deg,T ernary*_ = 0*/*h. Simulations were performed using all other parameters listed in Tables 3 and 4. Controlling for permeability, ACBI1 induces more efficient target degradation than PROTAC 1*.

**Figure 4:**
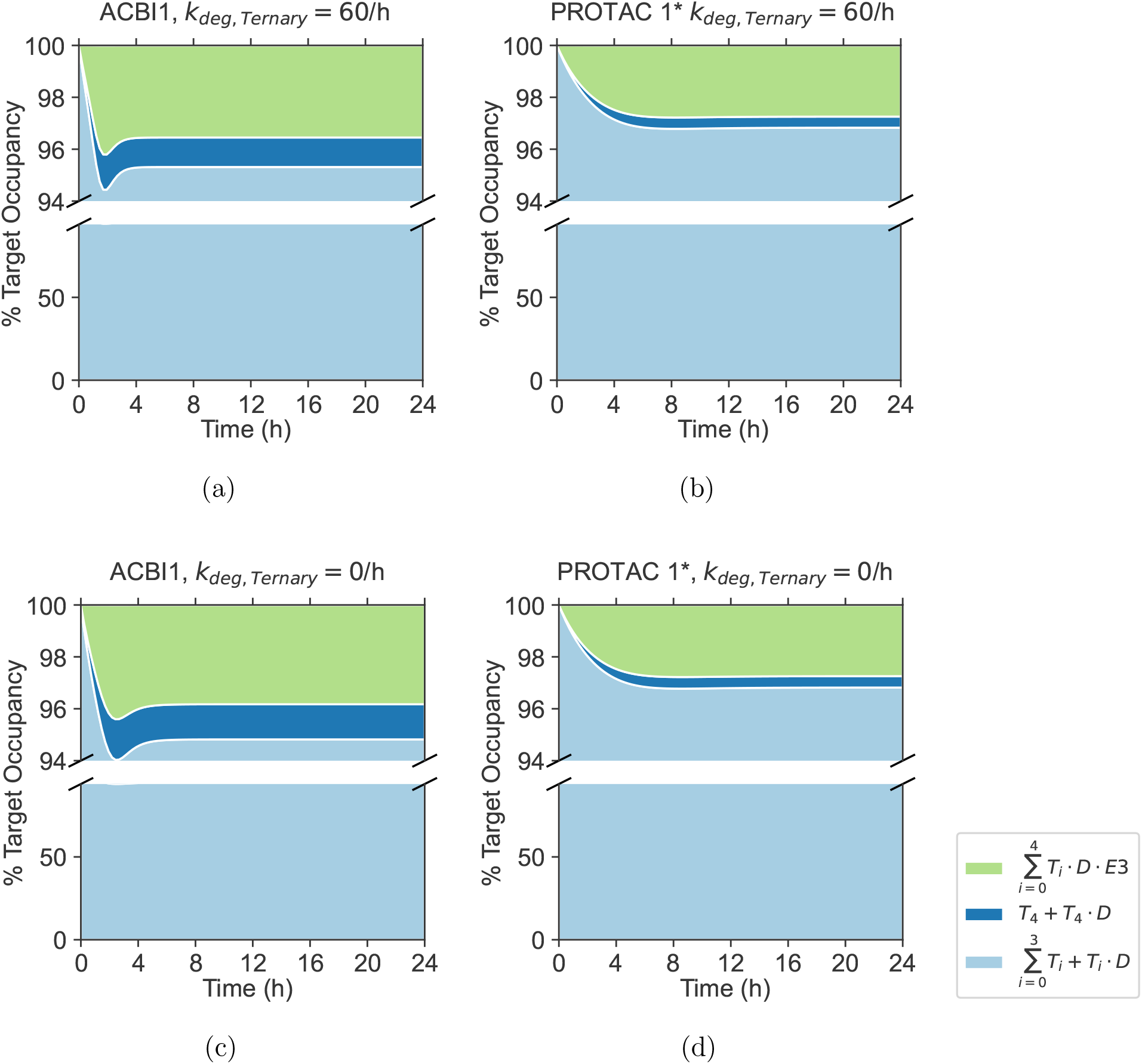
Percentage populations of target protein molecular species over time. Categories of species include insufficiently ubiquitinated free target and binary complex 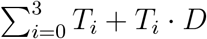, polyubiquitinated target molecules primed for degradation *T*_4_ + *T*_4_ · *D*, and total ternary Complex 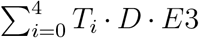. The relative population is calculated as the percentage of the total target concentration at each time point. Permeability for the PROTAC 1* molecule was set to the *PS*_*cell*_ parameter value for ACBI1 in Table 4. With different ternary complex formation kinetics and all other simulation parameters equal, a greater percentage of total target is polyubiquitinated or bound to a ternary complex under ACBI1-mediated degradation.

**Figure 5:**
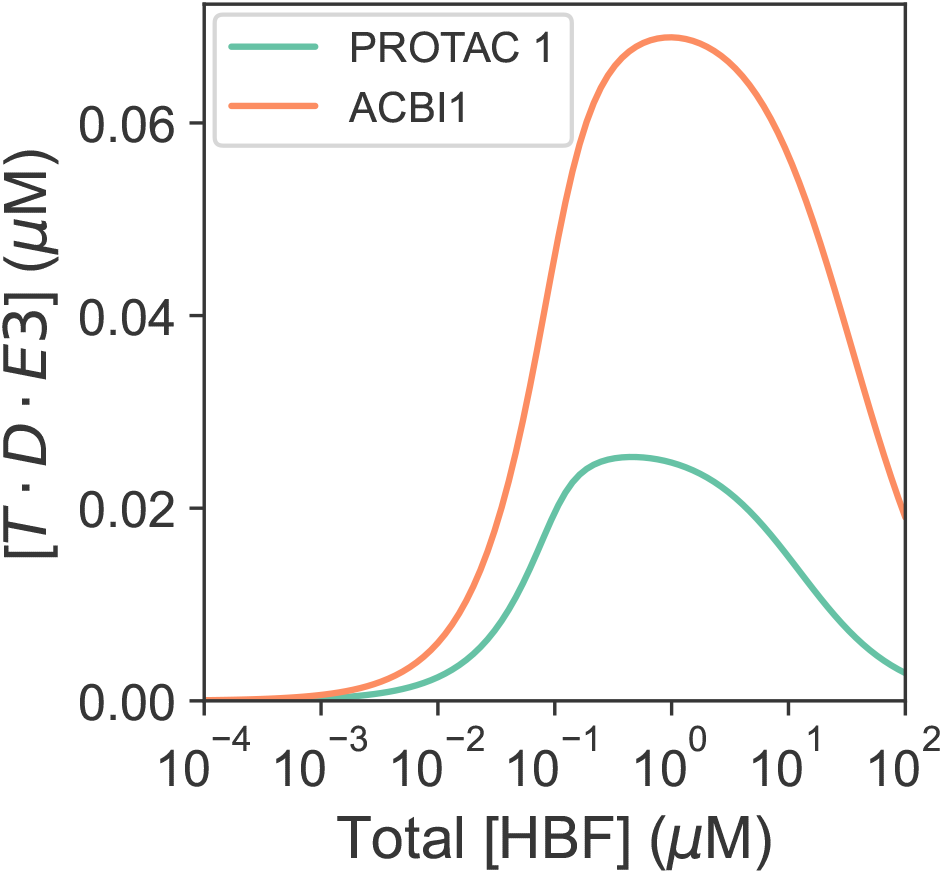
ACBI1- and PROTAC 1-induced ternary complex formation at equilibrium as a function of total intracellular degrader concentration. Total concentrations fixed for target at 1 *µ*M and for E3 at 0.1 *µ*M. Equilibrium solutions were calculated using molecule-specific binary binding affinities and cooperativity values listed in Table 4. Simulated ternary complex formation curves exhibit a hook effect.

Our model demonstrates that the kinetic process of degradation also exhibits this non-monotonous dependency on HBF concentration (Fig. 6). There are, however, important differences from the equilibrium hook effect. In our simulation of ACBI1, for example, the decrease in degradation at time *t* = 6 h at high degrader concentration is not associated with a decrease in the ternary complex formation. Also, degradation remains relatively constant over a wide range of degrader concentrations, wider in the case of ACBI1 (higher cooperativity) than in the case of PROTAC 1 (lower cooperativity). The dependency of ternary complex formation (at fixed time points) on the degrader concentration shows more complex behavior than equilibrium ternary complex formation, for the following reasons. In the degradation process, the formation of the ternary complex leads to target degradation, which creates a reactive flux that in effect suppresses the ternary complex concentration. As a result, the ternary complex concentration does not reach its equilibrium value, but stays relatively flat at a lower steady state value due to the balance between ternary complex formation and the subsequent target degradation, until the degrader concentration reaches a threshold where the equilibrium value of ternary complex concentration is approximately the same as the steady state value. Above this threshold the ternary complex concentration decreases with increasing degrader concentration. This threshold is reached in the case of PROTAC 1 but not ACBI1 in our simulations.

**Figure 6:**
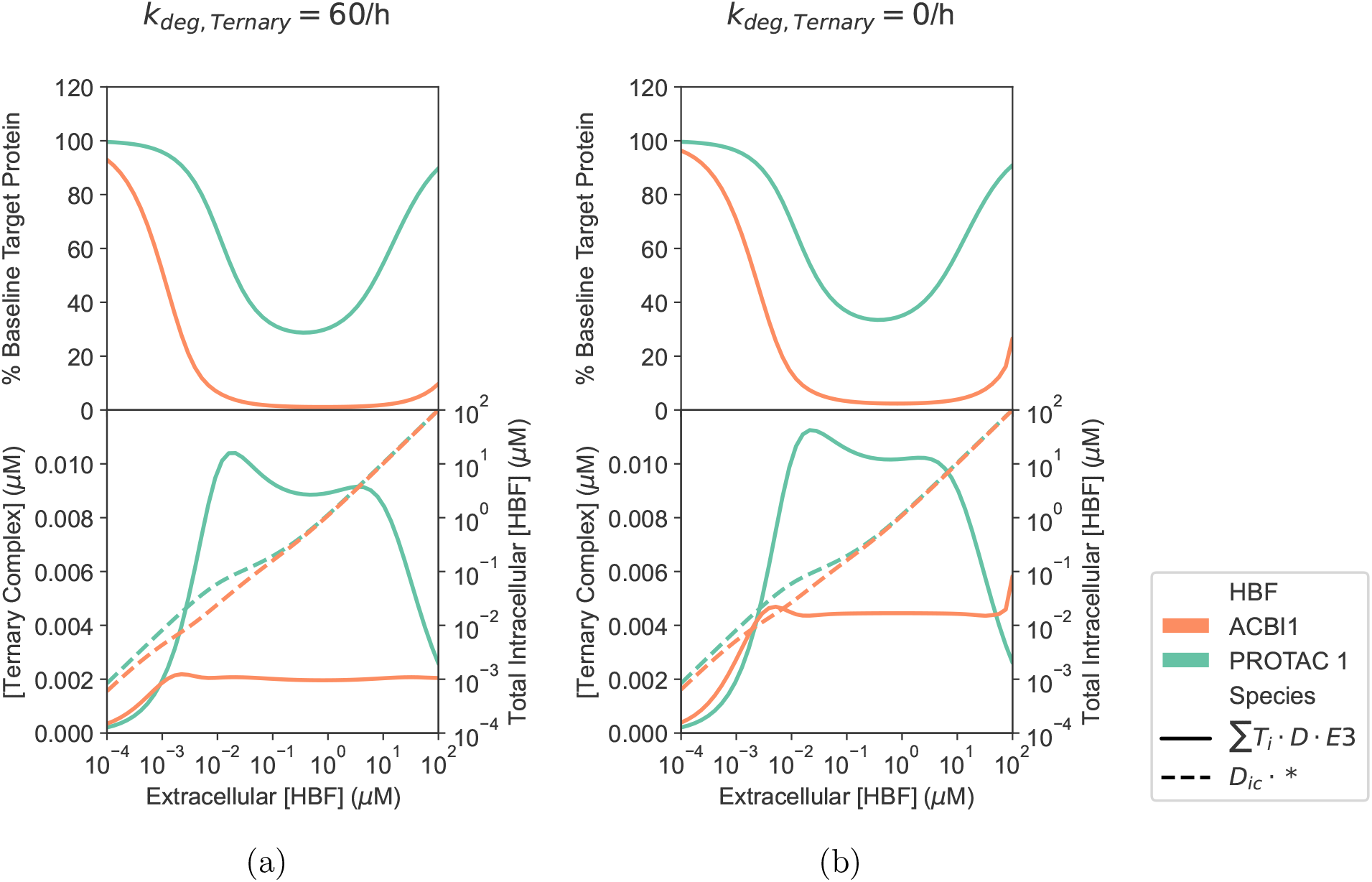
ACBI1- and PROTAC 1-mediated TPD; ternary complex formation, 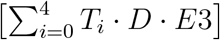; and total intracellular HBF concentration, [*D*_*ic*_ · *], at *t* = 6 h as a function of initial extracellular HBF exposure. Total E3 concentration was fixed at 0.1 *µ*M. Ternary complex degradation rate *k*_*deg,T ernary*_ was compared at two levels: baseline 0/h and equal to poly-ubiquitinated target degradation rate *k*_*deg,T*_ = 60*/*h. Simulations were performed with all other parameters fixed at values listed in Tables 3 and 4.

Next, we investigate the relationship between the stability of the ternary complex and degradation efficiency by exploring how changing the cooperativity in ternary complex formation affects degradation. We vary the value of cooperativity, *α*, by simultaneously multiplying *k*_*off,T,ternary*_, *K*_*D,T,ternary*_, *k*_*off,E3,ternary*_, and *K*_*D,E3,ternary*_ (in Table 4) by a scaling factor, keeping the other kinetic parameters constant. Unless otherwise noted, the parameters associated with ACBI1 are used to perform simulations.

If the polyubiquitinated target is degraded efficiently without first dissociating from the ternary complex, *e*.*g*. if *k*_*deg,T ernary*_ = *k*_*deg,T*_, target degradation increases with cooperativity, reaching saturation at different values of *α* for different binary affinities (Fig. 7a). When the HBF degrader has a strong binary affinity for both the target and the E3 ligase, efficient degradation takes place even with negative cooperativity (*α <* 1). If the binary affinity between the HBF and, for example, the target is weak, high cooperativity is required to form a stable ternary complex and to induce degradation. Our results explain the experimental observation that efficient HBF degraders may have a wide range of cooperativity values.^30^ In this scenario, highly cooperative—thus stable—ternary complex implies efficient degradation.

**Figure 7:**
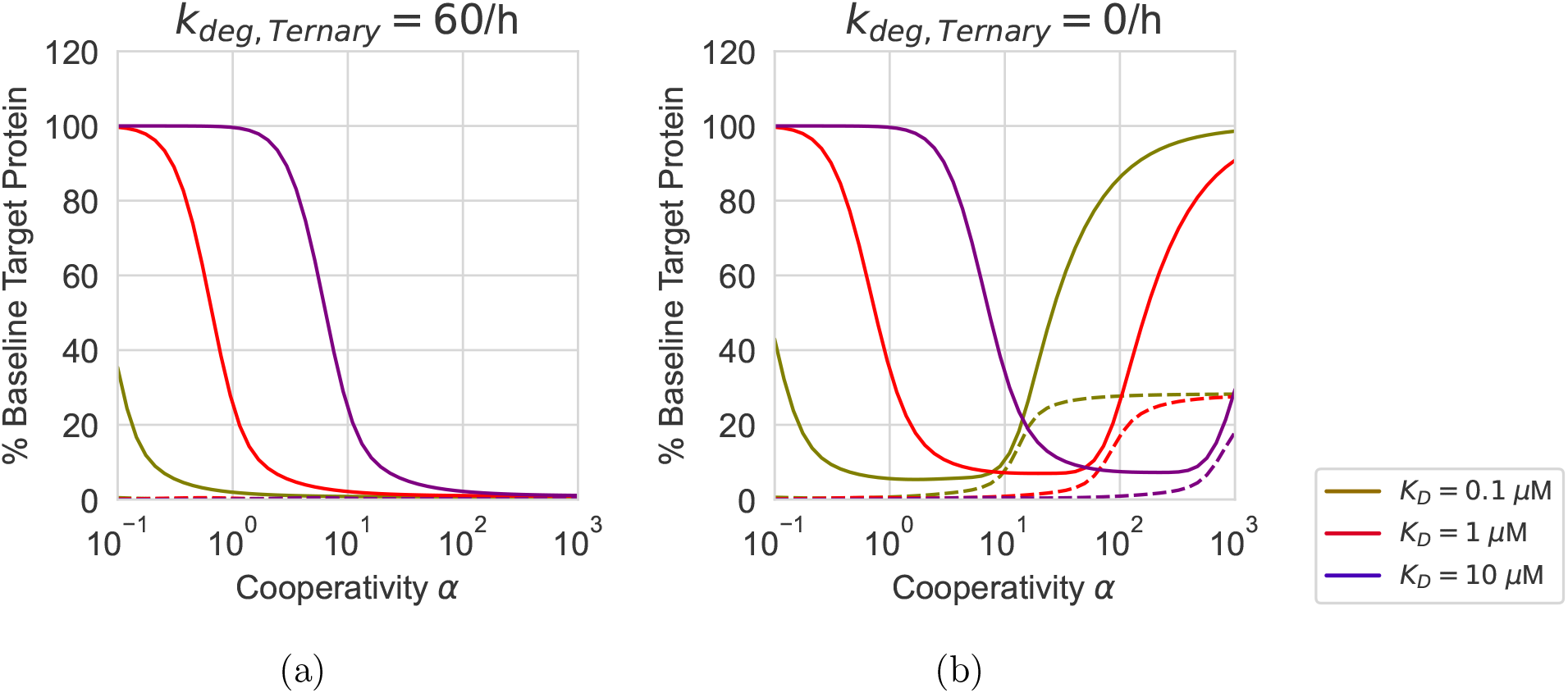
Simulated target protein degradation (solid lines) and ternary complex formation (dashed lines) at *t* = 6 h as a function of cooperativity and HBF-target binding affinity. *K*_*D,T,binary*_ values listed in the legend were implemented as a change in *k*_*off,T,binary*_. Ternary complex degradation rate *k*_*deg,T ernary*_ was compared at two levels: baseline 0/h and equal to poly-ubiquitinated target degradation rate *k*_*deg,T*_ = 60*/*h. The threshold of cooperativity past which degradation starts to diminish increases with *K*_*D,T,binary*_ when *k*_*deg,T ernary*_ = 0*/*h.

At the opposite limit that the polyubiquitinated target in the ternary complex cannot be degraded by the ternary complex, *i*.*e. k*_*deg,T eranry*_ = 0, we find that there is a non-monotonous relationship between target protein degradation and cooperativity, and the range of cooperativity values associated with efficient degradation depends on the binding constants of the binary HBF-target (or HBF-E3) complex (Fig. 7b). Our model predicts diminished degradation when cooperativity exceeds a certain threshold, the value of which increases with the value of the binary *K*_*D*_. At high cooperativity, the polyubiquitinated target dissociates slowly from the highly stable ternary complex, which, under the condition of *k*_*deg,T ernary*_ ≈ 0, protects the target from degradation. Our analysis thus suggests that under certain conditions there may be an optimal range of cooperativity for efficient degradation, and that this range depends on the binary affinity between the HBF and the target (and the binary affinity between the HBF and the E3 ligase). This may help explain the experimental observation that cooperative ternary complex does not necessarily lead to efficient degradation.^7,30,31^

Given our model’s novel prediction of an optimal range of cooperativity values for degradation under the limiting scenario of *k*_*deg,T ernary*_ = 0, we next explore the conditions—under this same limiting scenario—that admit the above non-monotonous relationship between cooperativity and degradation, and how the optimal range of cooperativity depends on the other kinetic parameters of the degradation process. We will then investigate the range of *k*_*deg,T ernary*_ within which degradation may decrease with increasing cooperativity.

Our model suggests that the optimal range of cooperativity depends on the ubiquitination rate (Fig. 8a). The target in the ternary complexes of different HBF molecules may have different ubiquitination rates because of the difference in their structural ensembles,^32,33^ which affects the exposure of lysines on the target surface to ubiquitination.^34^ If ubiquitination is fast, a transient ternary complex is sufficient for ubiquitination to take place, and fast dissociation of the ternary complex—corresponding to a low cooperativity—leads to both high turnover of the HBF molecules and rapid production of free, polyubiquitinated target ready for degradation; in contrast, high cooperativity—and thus a very stable ternary complex—may hinder the catalytic turnover and slow down the degradation.

**Figure 8:**
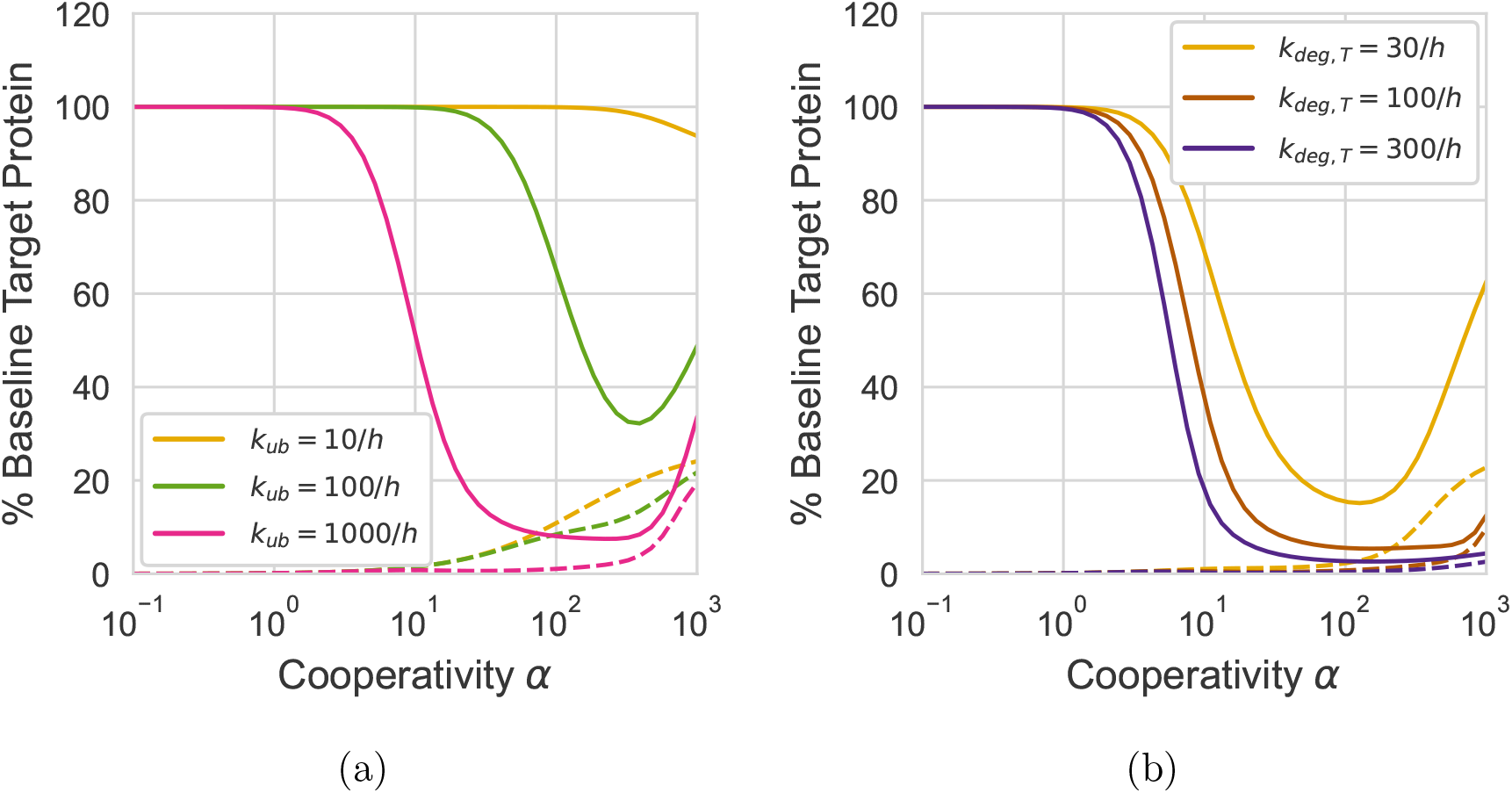
Simulated target protein degradation (solid lines) and ternary complex formation (dashed lines) at *t* = 6 h as a function of cooperativity and: (a) ubiquitination rate (*k*_*ub*_ in Table. 3), (b) proteasomal degradation rate (*k*_*deg,T*_ in Table. 3) with *k*_*ub*_ fixed at 1000*/h*.

Our model predicts that non-monotonous dependence of degradation on cooperativity only holds if the degradation of polyubiquitinated target is slow. If this degradation step is sufficiently fast, efficient degradation occurs despite slow dissociation of polyubiquitinated target from a highly stable ternary complex (Fig. 8b).

Different target proteins have different baseline cellular concentrations, which affect their degradability. Here we investigate how the optimal range of cooperativity depends on a target’s baseline cellular concentration (Fig. 9). For targets with low cellular concentrations, efficient degradation can occur over a wide range of cooperativity values. For targets with high cellular concentrations, cooperativity needs to fall inside a very narrow range to enable efficient degradation, because it requires high catalytic turnover to degrade a large quantity of target proteins.

**Figure 9:**
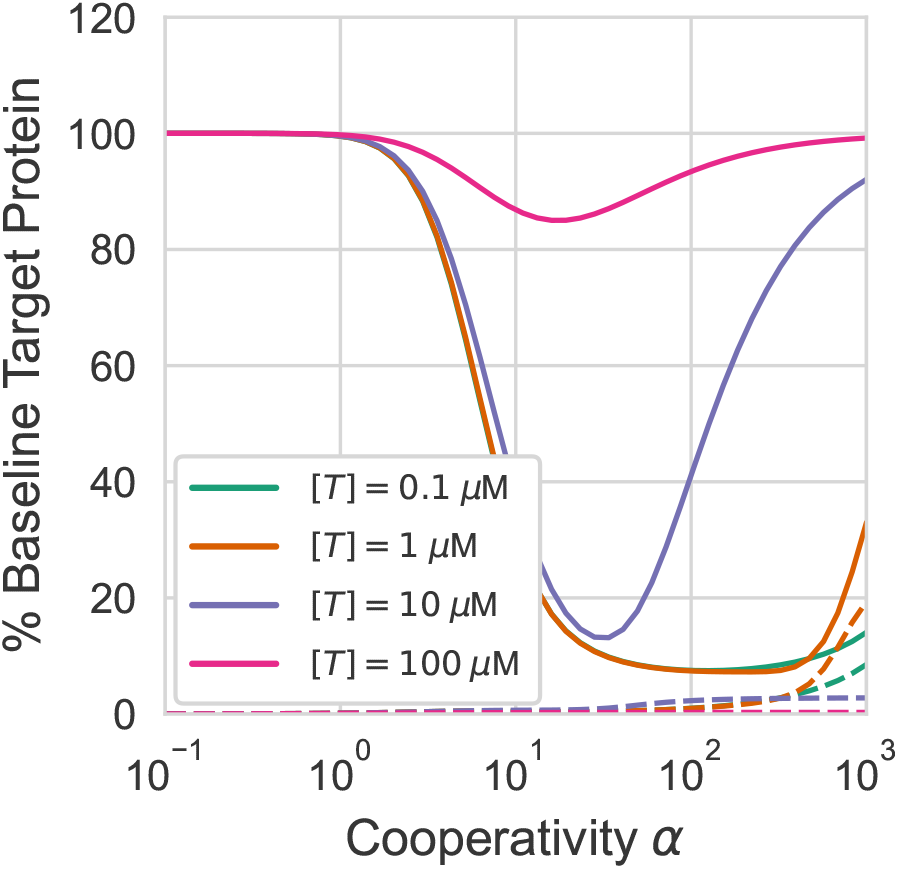
Simulated target protein degradation (solid lines) and ternary complex formation (dashed lines) at *t* = 6 h as a function of cooperativity and baseline target POI expression. Baseline target concentration was changed by changing the target production rate (*k*_*prod*_ in Table 3) according to Eq. 1. The range of cooperativity values that induce efficient degradation becomes more restricted as baseline target concentration increases.

We have primarily analyzed the effect of cooperativity on degradation at an intermediate time *t* = 6h. Here we note that the effect depends on the time points of measured degradation (Fig. 10a). There is a narrower range of cooperativity for effective degradation at short times (*e*.*g. t* = 1h) than at longer times (*e*.*g. t* = 6, 12h). At short times, degradation mediated by an HBF with low cooperativity is limited by ternary complex formation (Fig. 4d; at *t* = 1h only 1.04% of the target is in the ternary complex for PROTAC 1 with *α* = 3.2), whereas degradation mediated by an HBF with high cooperativity (*e*.*g*., *α* = 200) is limited by the dissociation of the polyubiquitinated target from the ternary complex to become proteasome degradable (Fig. 10b; at *t* = 1h only 0.4% of the target is polyubiquitinated and not bound in a ternary complex). As a result, effective degradation requires a narrow range of cooperativity. At longer times, in contrast, the system is in a steady state with a sufficient accumulation of free, polyubiquitinated target protein to sustain efficient degradation, even when the high cooperativity leads to a predominant population of ternary complex (at *t* = 6h, for an HBF with *α* = 200, even though 23% of the target is in a ternary complex, 2% of the target is polyubiquitinated and free). Thus efficient degradation still occurs at high *α* values (Fig. 10a).

**Figure 10:**
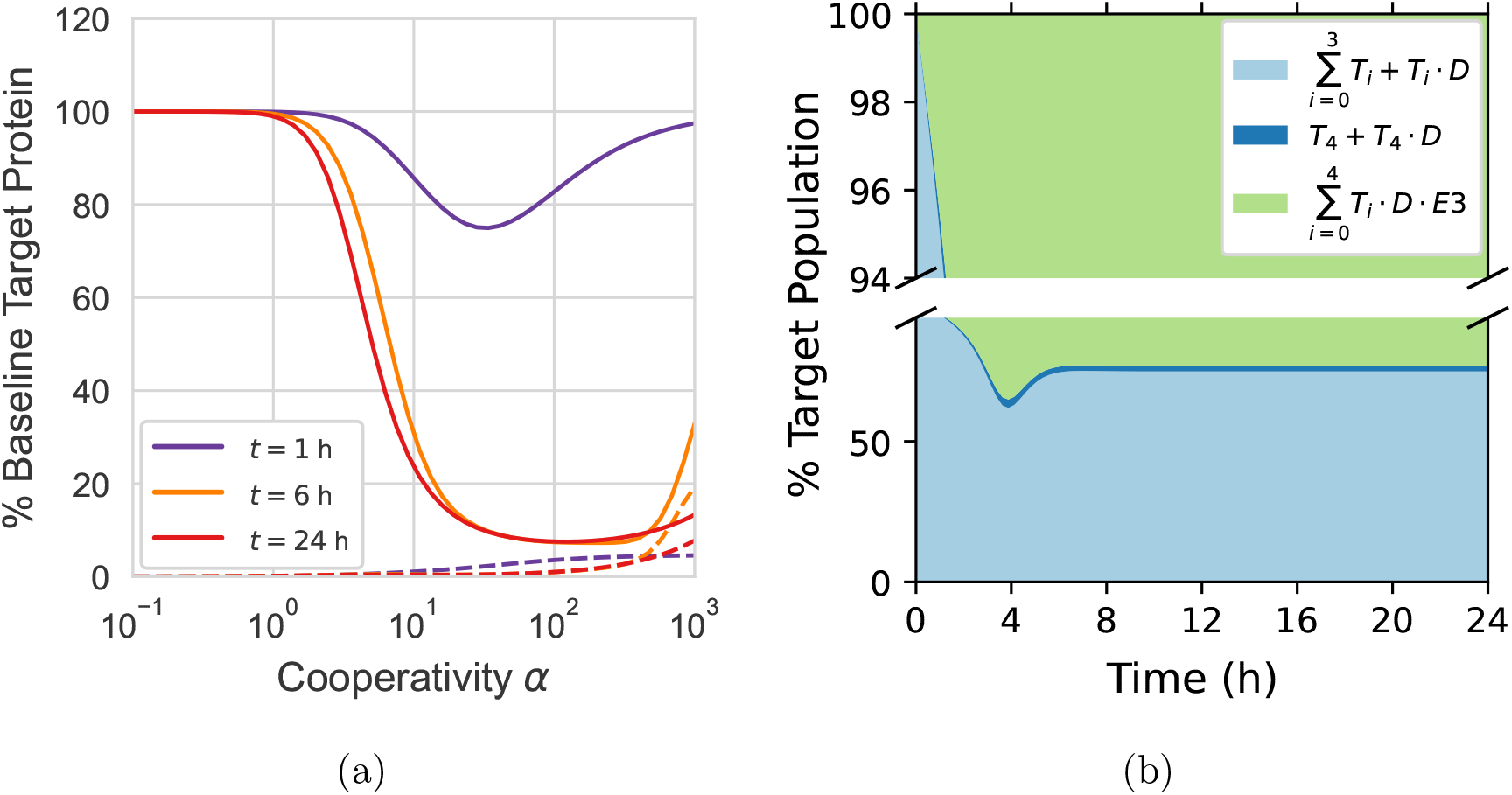
(a) Simulated target protein degradation (solid lines) and ternary complex formation (dashed lines) as a function of cooperativity and time. (b) Percentage populations of target protein molecular species over time. Categories of species include insufficiently ubiquitinated free target and binary complex 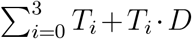, polyubiquitinated target molecules primed for degradation *T*_4_ + *T*_4_ · *D*, and total ternary complex 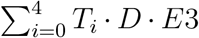. The relative population is calculated as the percentage of the total target concentration at each time point. Cooperativity was fixed at *α* = 200.

The above analyses considered only the limiting case of *k*_*deg,T ernary*_ = 0. A more realistic scenario may be 0 *< k*_*deg,T ernary*_ *< k*_*deg,T*_ (see Discussion). Fig. 11 shows how degradation depends on cooperativity at different values of *k*_*deg,T ernary*_. The non-monotonous relationship between degradation and cooperativity is only manifest when *k*_*deg,T ernary*_ is at least an order-of-magnitude slower than *k*_*deg,T*_.

**Figure 11:**
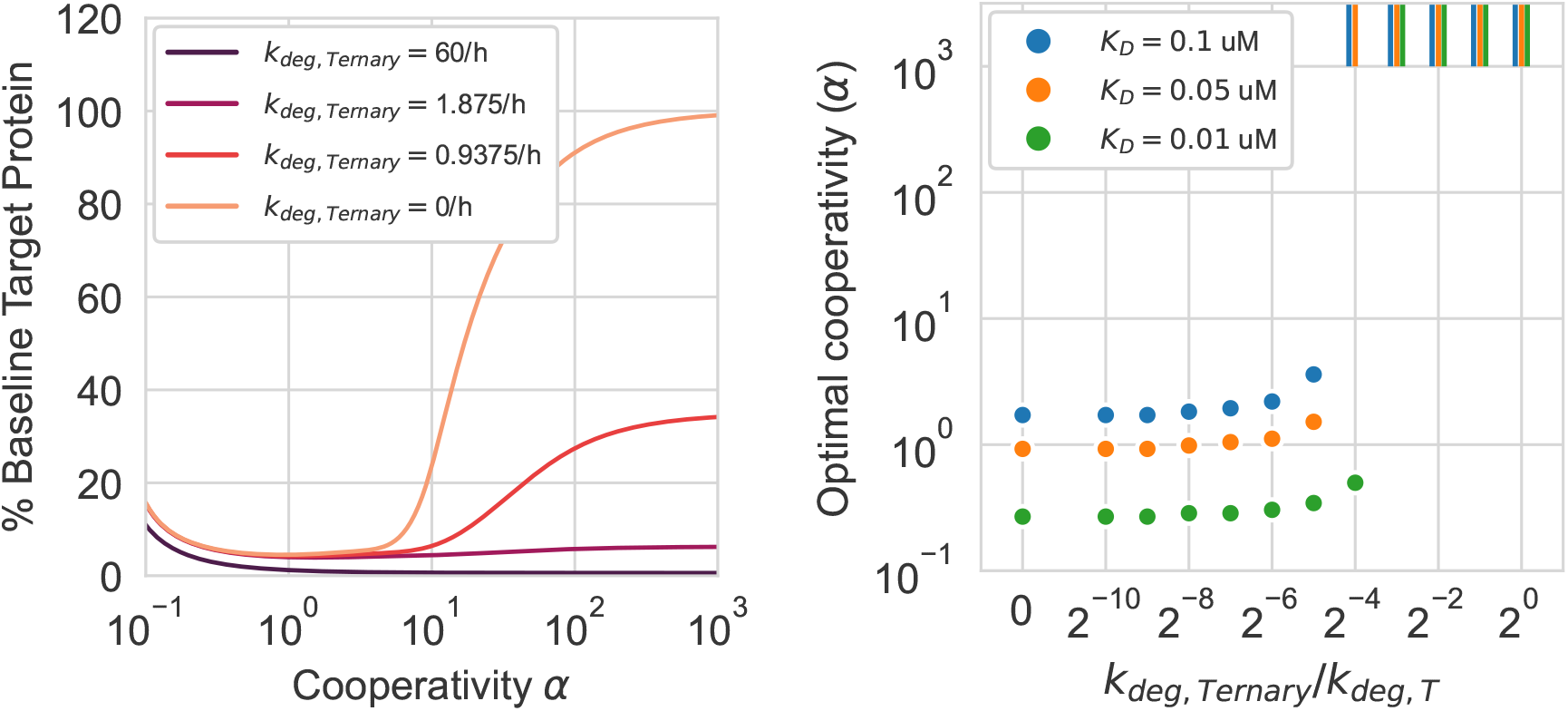
(a) Simulated target protein degradation at *t* = 6 h as a function of cooperativity and poly-ubiquitinated ternary complex degradation rate *k*_*deg,T ernary*_. Binary HBF-target *K*_*D*_ was fixed at 0.05 *µ*M. (b) Optimal cooperativity—the *α* value at which degradation at *t* = 6 h reaches a maximum—as a function of the ratio *k*_*deg,T ernary*_*/k*_*deg,T*_ across different values of binary target *K*_*D*_ with *k*_*deg,T*_ fixed at 60/h. Vertical bars indicate that the optimal cooperativity lies beyond the greatest *α* (10^3^) used in degradation simulations.

### Understanding the contributions of PPI to cooperativity

PPI between the target protein and the E3 ligase may affect the stability of the ternary complex. We hypothesized that introducing mutations at the target-E3 interface by protein site-directed mutagenesis could alter cooperativity. To test this hypothesis, we leveraged our statistical inference model developed in this work (see Methods) to analyze NanoBRET assays that measure intracellular ternary complex formation of SMARCA2-ACBI1-VHL at a range of ACBI1 concentrations, for wild-type (WT) SMARCA2 and VHL, two SMARCA2 mutants: LEU1465→SER (L1465S) and GLU1420→SER (E1420S), and two VHL mutants: TYR112→PHE (Y112F) and ARG69→GLN (R69Q). Based on the crystal structure of the SMARCA2-ACBI1-VHL ternary complex (Fig. 12a), the L1465S mutation in SMARCA2 and the Y112F and R69Q mutations in VHL are hypothesized to reduce the ternary complex stability, while the mutation E1420S should not. In this work, we demonstrate our inference model by providing a quick but approximate estimate of cooperativity of ternary complex formation in the cell, by making two simplifying assumptions that the mutations do not affect the stability and expression levels of SMARCA2 and VHL or their binary affinities for ACBI1. In a more detailed analysis, these changes in the expression levels of the target and the E3 ligase can be measured for the mutants, and the binary affinities can be measured using purified proteins, and these measured changes can be incorporated into our model to improve the accuracy of the inferred values of cooperativity.

**Figure 12:**
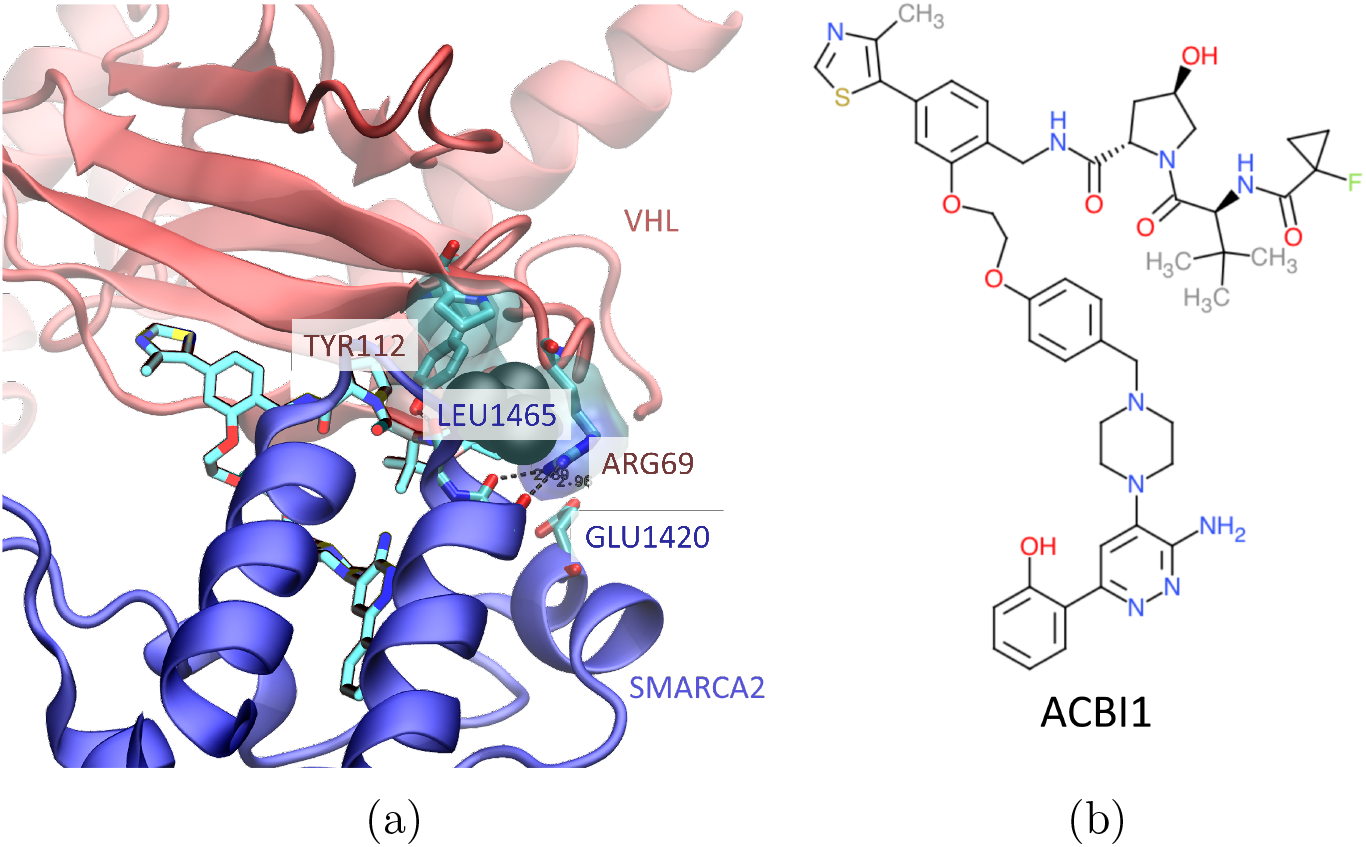
Probing the contribution of protein-protein interactions to cooperativity. (a) Protein-protein interactions in the SMARCA2-ACBI1-VHL ternary complex (PDB ID: 6HAY). TYR112 and ARG69 of VHL and LEU1465 of SMARCA2 form key interactions at the protein-protein interface. GLU1420 of SMARCA2 is solvent exposed and selected for control mutation. (b) Molecular structure of ACBI1.

Our inference model applied to the NanoBRET experimental data allows us to quantify the contributions of specific PPI to the cooperativity. Our model reasonably reproduces the titration curve observed in the experiments; both modeled and observed curves demonstrate the hook effect (Fig 13a). The mutations that abolish crystallographically observed protein-protein interactions—L1465S in SMARCA2, Y112F and R69Q in VHL—indeed reduce the cooperativity: posterior means 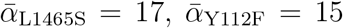, and 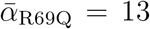, compared to *α*_WT_ = 26, which is fixed in our analysis to the value previously determined by surface plasmon resonance (unpublished results; the cooperativity was previously measured to be 28 by fluorescence polarization and 30 by time-resolved fluorescence energy transfer^6^). The above-mentioned simplifications and the difference between in-cell and biochemical assays may contribute to the quantitative difference between our result (*α* = 13) and the previously published *α* value (3.3)^6^ for the VHL:R69Q mutant. The posterior parameter distributions approximated by MCMC sampling exhibit a clear separation between these mutants and the wild-type: at least 97.5% of their probability densities lie below *α*_WT_ (Fig 13b). In contrast, the posterior distribution for the cooperativity parameter of the control mutant E1420S is centered at the posterior mean 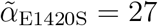, near the value of *α*_WT_.

**Figure 13:**
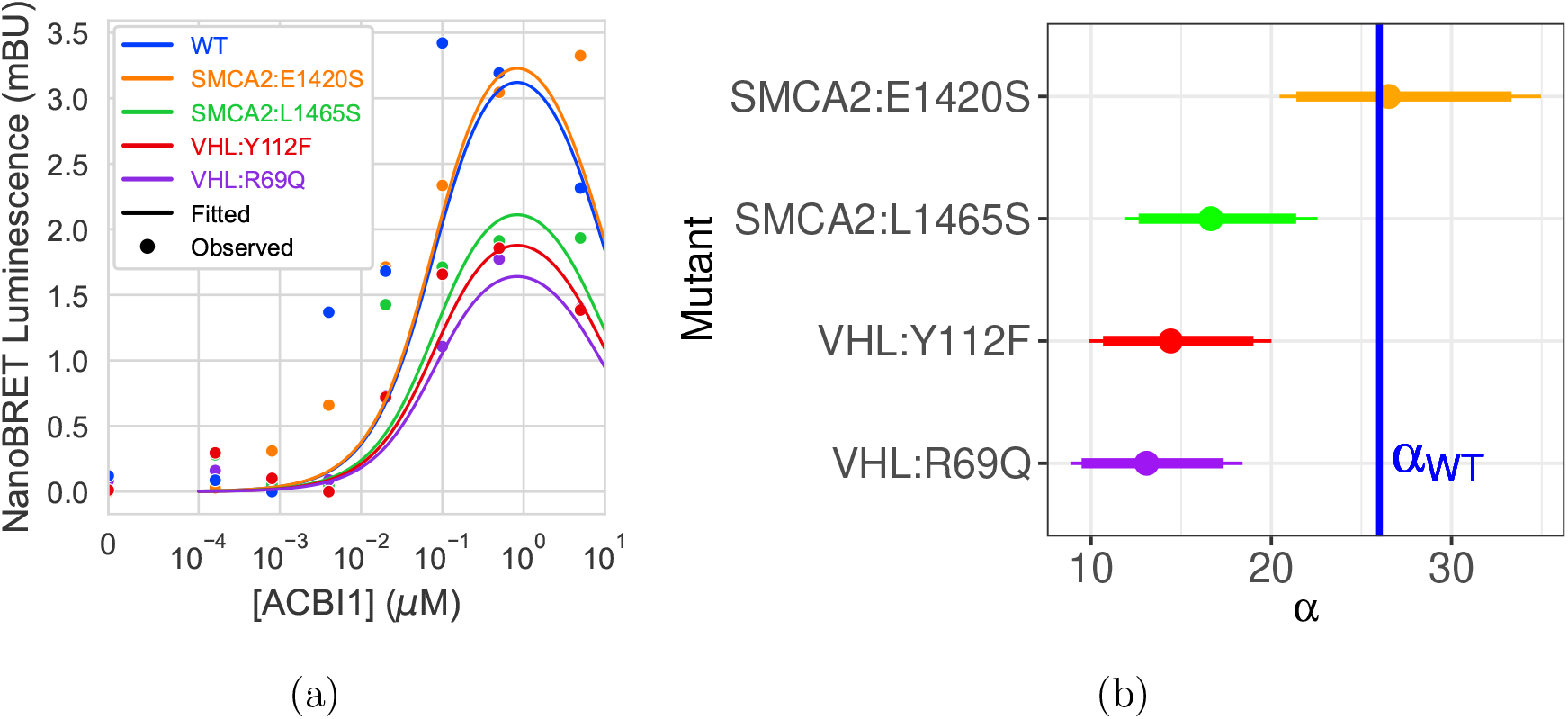
(a) Fitted versus observed ternary complex expression curves. Fitted curves were calculated using the posterior mean mutant-specific cooperativity and values for all other parameters sampled from the posterior distribution. (b) Posterior median and highest posterior density intervals for cooperativity parameters labeled by protein mutation. Thick lines = 90% posterior density intervals, thin lines = 95%. Wild type cooperativity *α*_WT_ = 26 shown in blue. Parameter values sampled by MCMC from the posterior distribution support the hypothesis that the mutations L1465S, Y112F, R69Q weaken ternary complex stability while the mutation E1420S does not.

## Discussion

This work focuses on relating the stability of the ternary complex to the degradation of the target protein. Accordingly, we have simplified several complex processes of multiple reactions subsequent to ternary complex formation into single-step reactions. For example, the degradation of the polyubiquitinated target, either free or bound in the ternary complex, requires first its binding to the proteasome particle (sometimes facilitated by the shuttle factors) and then the deubiquitination and unfolding of the target protein by an energy-consuming, active process on the proteasome.^35,36^ The unfolded single-chain protein is then translocated through a ringed channel and degraded.^37^ Each is a separate step with its own rate of reaction. The rate of binding to the proteasome depends on the diffusivity of the molecular species (the ternary complex may diffuse slower than the free target protein), the extent of ubiquitination, and the proteasome concentration at the cellular locus. The rate of deubiquitination and unfolding may depend on the stability of the target protein, its extent of ubiquitination, and the presence or absence of other bound proteins, including the E3 ligase in the ternary complex (the complex may stabilize the target protein and retard its deubiquitination and unfolding). Including these steps in detail in our model would result in a complex system of reactions with too many unknown parameters. Instead, we have reduced the degradation of the polyubiquitinated target into a single step, with one rate for the free target (*k*_*deg,T*_) and another for the target in the ternary complex (*k*_*deg,T ernary*_), assuming *k*_*deg,T ernary*_ ≤ *k*_*deg,T*_ for the above reasons.

Despite these simplifications, our model reveals the relationship between degradation and ternary complex stability under different conditions defined by kinetic rates of the key steps of TPD. Extending the previous pharmacodynamic models ^22,23^ of TPD to explicitly include the deubiquitination cascade and different degradation rates for the free ubiquitinated target and the target in the ternary complex, our model enables us to investigate the consequences of these parameters on the degradation and make novel predictions, such as the possibility of an optimal range of cooperativity for degradation under certain conditions. Our model may help determine when modulating the ternary complex stability will substantially impact degradation, and in which direction.

The unique prediction of our model—that an over-stabilized ternary complex may hinder degradation—requires that a free polyubiquitinated target protein is degraded much faster than one in a ternary complex, *i*.*e. k*_*deg,T*_ ≫ *k*_*deg,T ernary*_. This prediction is yet to be validated by experimental examples that satisfy the conditions underlying the prediction. Because the optimal range of cooperativity depends on many kinetic parameters, a highly stable ternary complex may still admit efficient degradation.^38,39^ To our knowledge, there have been no reports of measured degradation kinetics subsequent to target polyubiquitination, let alone separate degradation kinetics of the free polyubiquitinated target and the ternary complex, leaving their relative rates an open question.

Many of the kinetic parameters in the cellular degradation process—needed in our model— have not been experimentally determined. We have guess-estimated their values in this work. For example, we estimated the degradation rate of a polyubiquitinated target, *k*_*deg,T*_, based on the experimental observation that degradation of a single chain of 300 amino acid residues takes more than 20 seconds after the substrate is bound to the proteasome;^36,40^ correspondingly a substrate such as SMARCA2 (1590 amino acids) may take up to 100 seconds. The true degradation rate and other kinetic parameters such as the ubiquitination rate *k*_*ub*_ and the deubiquitination rate *k*_*deub*_ depend on the cellular context including the concentrations of the proteasome, E2 enzymes, and deubiquitinases. These kinetic parameters may be determined by fitting our model to an experimentally observed time course of HBF-induced degradation.^21^

## Conclusions

Our pharmacodynamic model of HBF-induced TPD enables us to quantitatively investigate how changes in the kinetic rates of the key steps in the degradation process affect the degradation efficiency. The development of therapeutic HBF degraders to degrade a POI entails many iterations of medicinal chemistry modifications of candidate molecules to improve their cellular and *in vivo* degradation efficacy. Such modifications change the stability of the ternary complex—including by way of changing the cooperativity—and the ubiquitination rate of the target by altering the structure and dynamics of the ternary complex and thus the accessibility of lysines on the POI’s surface. Our model may help identify which changing parameters are driving the changes in degradation efficiency, thus guiding the optimization of HBF degraders in the relevant dimensions and in the correct directions. Our model can estimate how degradation efficiency depends on other parameters, such as the baseline concentration of the target POI, and thus it may inform the feasibility assessment of a POI as a target for therapeutic intervention by degradation. In addition, our mathematical framework may be extended to analyze the catalytic efficiency of other types of chemically induced proximity phenomena.^41,42^

## Data and Software Availability

The Python code for the pharmacodynamic model and the R script for the statistical inference model will be publicly available in GitHub upon manuscript publication. The experimental data in Fig. 13a will be deposited with the R script.

## Appendix

## Acknowledgement

We thank Woody Sherman and Frances Rodriguez-Rivera for discussions and suggestions to the manuscript and Yunxing (Stella) Li for preparing the Table of Content (TOC) graphic.

## Supporting Information Available

The nucleic acid sequences of the cloned vectors.

## TOC Graphic

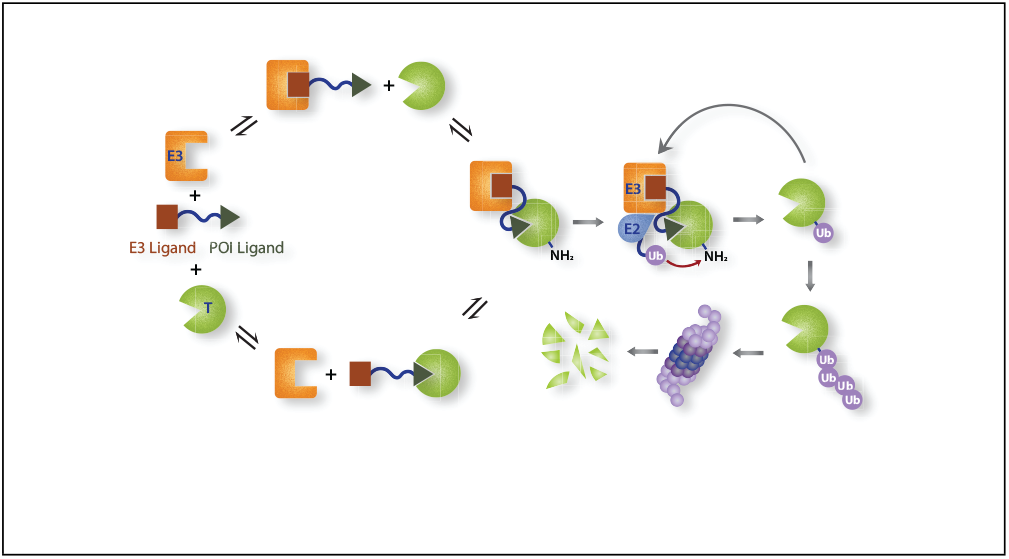

