## Supplementary Information for "Modeling the effect of cooperativity in ternary complex formation and targeted protein degradation mediated by heterobifunctional degraders"

**Halo-VHL and nLuc-SMARCA2 sequences**

nLuc-SMARCA2 includes amino acids 1373-1511 representing the same amino acids within 6HAY PDB.

Halo-VHL

ATGGCAGAAATCGGTACTGGCTTTCCATTCGACCCCCATTATGTGGAAGTCCTGGGCGAGCGCATGCACTACGTCGATGTTGGTCCGCGCGATGGCACCCCTGTGCTGTTCCTGCACGGTAACCCGACCTCCTCCTACGTGTGGCGCAACATCATCCCGCATGTTGCACCGACCCATCGCTGCATTGCTCCAGACCTGATCGGTATGGGCAAATCCGACAAACCAGACCTGGGTTATTTCTTCGACGACCACGTCCGCTTCATGGATGCCTTCATCGAAGCCCTGGGTCTGGAAGAGGTCGTCCTGGTCATTCACGACTGGGGCTCCGCTCTGGGTTTCCACTGGGCCAAGCGCAATCCAGAGCGCGTCAAAGGTATTGCATTTATGGAGTTCATCCGCCCTATCCCGACCTGGGACGAATGGCCAGAATTTGCCCGCGAGACCTTCCAGGCCTTCCGCACCACCGACGTCGGCCGCAAGCTGATCATCGATCAGAACGTTTTTATCGAGGGTACGCTGCCGATGGGTGTCGTCCGCCCGCTGACTGAAGTCGAGATGGACCATTACCGCGAGCCGTTCCTGAATCCTGTTGACCGCGAGCCACTGTGGCGCTTCCCAAACGAGCTGCCAATCGCCGGTGAGCCAGCGAACATCGTCGCGCTGGTCGAAGAATACATGGACTGGCTGCACCAGTCCCCTGTCCCGAAGCTGCTGTTCTGGGGCACCCCAGGCGTTCTGATCCCACCGGCCGAAGCCGCTCGCCTGGCCAAAAGCCTGCCTAACTGCAAGGCTGTGGACATCGGCCCGGGTCTGAATCTGCTGCAAGAAGACAACCCGGACCTGATCGGCAGCGAGATCGCGCGCTGGCTGTCGACGCTCGAGATTTCCGGCGAGCCAACCACTGAGGATCTGTACTTTCAGAGCGATAACGCGATCGCCATGCCCCGGAGGGCGGAGAACTGGGACGAGGCCGAGGTAGGCGCGGAGGAGGCAGGCGTCGAAGAGTACGGCCCTGAAGAAGACGGCGGGGAGGAGTCGGGCGCCGAGGAGTCCGGCCCGGAAGAGTCCGGCCCGGAGGAACTGGGCGCCGAGGAGGAGATGGAGGCCGGGCGGCCGCGGCCCGTGCTGCGCTCGGTGAACTCGCGCGAGCCCTCCCAGGTCATCTTCTGCAATCGCAGTCCGCGCGTCGTGCTGCCCGTATGGCTCAACTTCGACGGCGAGCCGCAGCCCTACCCAACGCTGCCGCCTGGCACGGGCCGCCGCATCCACAGCTACCGAGGTCACCTTTGGCTCTTCAGAGATGCAGGGACACACGATGGGCTTCTGGTTAACCAAACTGAATTATTTGTGCCATCTCTCAATGTTGACGGACAGCCTATTTTTGCCAATATCACACTGCCAGTGTATACTCTGAAAGAGCGATGCCTCCAGGTTGTCCGGAGCCTAGTCAAGCCTGAGAATTACAGGAGACTGGACATCGTCAGGTCGCTCTACGAAGATCTGGAAGACCACCCAAATGTGCAGAAAGACCTGGAGCGGCTGACACAGGAGCGCATTGCACATCAACGGATGGGAGATGTTTAA

Halo-VHL R69Q

ATGGCAGAAATCGGTACTGGCTTTCCATTCGACCCCCATTATGTGGAAGTCCTGGGCGAGCGCATGCACTACGTCGATGTTGGTCCGCGCGATGGCACCCCTGTGCTGTTCCTGCACGGTAACCCGACCTCCTCCTACGTGTGGCGCAACATCATCCCGCATGTTGCACCGACCCATCGCTGCATTGCTCCAGACCTGATCGGTATGGGCAAATCCGACAAACCAGACCTGGGTTATTTCTTCGACGACCACGTCCGCTTCATGGATGCCTTCATCGAAGCCCTGGGTCTGGAAGAGGTCGTCCTGGTCATTCACGACTGGGGCTCCGCTCTGGGTTTCCACTGGGCCAAGCGCAATCCAGAGCGCGTCAAAGGTATTGCATTTATGGAGTTCATCCGCCCTATCCCGACCTGGGACGAATGGCCAGAATTTGCCCGCGAGACCTTCCAGGCCTTCCGCACCACCGACGTCGGCCGCAAGCTGATCATCGATCAGAACGTTTTTATCGAGGGTACGCTGCCGATGGGTGTCGTCCGCCCGCTGACTGAAGTCGAGATGGACCATTACCGCGAGCCGTTCCTGAATCCTGTTGACCGCGAGCCACTGTGGCGCTTCCCAAACGAGCTGCCAATCGCCGGTGAGCCAGCGAACATCGTCGCGCTGGTCGAAGAATACATGGACTGGCTGCACCAGTCCCCTGTCCCGAAGCTGCTGTTCTGGGGCACCCCAGGCGTTCTGATCCCACCGGCCGAAGCCGCTCGCCTGGCCAAAAGCCTGCCTAACTGCAAGGCTGTGGACATCGGCCCGGGTCTGAATCTGCTGCAAGAAGACAACCCGGACCTGATCGGCAGCGAGATCGCGCGCTGGCTGTCGACGCTCGAGATTTCCGGCGAGCCAACCACTGAGGATCTGTACTTTCAGAGCGATAACGCGATCGCCATGCCCCGGAGGGCGGAGAACTGGGACGAGGCCGAGGTAGGCGCGGAGGAGGCAGGCGTCGAAGAGTACGGCCCTGAAGAAGACGGCGGGGAGGAGTCGGGCGCCGAGGAGTCCGGCCCGGAAGAGTCCGGCCCGGAGGAACTGGGCGCCGAGGAGGAGATGGAGGCCGGGCGGCCGCGGCCCGTGCTGCGCTCGGTGAACTCGCAGGAGCCCTCCCAGGTCATCTTCTGCAATCGCAGTCCGCGCGTCGTGCTGCCCGTATGGCTCAACTTCGACGGCGAGCCGCAGCCCTACCCAACGCTGCCGCCTGGCACGGGCCGCCGCATCCACAGCTACCGAGGTCACCTTTGGCTCTTCAGAGATGCAGGGACACACGATGGGCTTCTGGTTAACCAAACTGAATTATTTGTGCCATCTCTCAATGTTGACGGACAGCCTATTTTTGCCAATATCACACTGCCAGTGTATACTCTGAAAGAGCGATGCCTCCAGGTTGTCCGGAGCCTAGTCAAGCCTGAGAATTACAGGAGACTGGACATCGTCAGGTCGCTCTACGAAGATCTGGAAGACCACCCAAATGTGCAGAAAGACCTGGAGCGGCTGACACAGGAGCGCATTGCACATCAACGGATGGGAGATGTTTAA

Halo-VHL Y112F

ATGGCAGAAATCGGTACTGGCTTTCCATTCGACCCCCATTATGTGGAAGTCCTGGGCGAGCGCATGCACTACGTCGATGTTGGTCCGCGCGATGGCACCCCTGTGCTGTTCCTGCACGGTAACCCGACCTCCTCCTACGTGTGGCGCAACATCATCCCGCATGTTGCACCGACCCATCGCTGCATTGCTCCAGACCTGATCGGTATGGGCAAATCCGACAAACCAGACCTGGGTTATTTCTTCGACGACCACGTCCGCTTCATGGATGCCTTCATCGAAGCCCTGGGTCTGGAAGAGGTCGTCCTGGTCATTCACGACTGGGGCTCCGCTCTGGGTTTCCACTGGGCCAAGCGCAATCCAGAGCGCGTCAAAGGTATTGCATTTATGGAGTTCATCCGCCCTATCCCGACCTGGGACGAATGGCCAGAATTTGCCCGCGAGACCTTCCAGGCCTTCCGCACCACCGACGTCGGCCGCAAGCTGATCATCGATCAGAACGTTTTTATCGAGGGTACGCTGCCGATGGGTGTCGTCCGCCCGCTGACTGAAGTCGAGATGGACCATTACCGCGAGCCGTTCCTGAATCCTGTTGACCGCGAGCCACTGTGGCGCTTCCCAAACGAGCTGCCAATCGCCGGTGAGCCAGCGAACATCGTCGCGCTGGTCGAAGAATACATGGACTGGCTGCACCAGTCCCCTGTCCCGAAGCTGCTGTTCTGGGGCACCCCAGGCGTTCTGATCCCACCGGCCGAAGCCGCTCGCCTGGCCAAAAGCCTGCCTAACTGCAAGGCTGTGGACATCGGCCCGGGTCTGAATCTGCTGCAAGAAGACAACCCGGACCTGATCGGCAGCGAGATCGCGCGCTGGCTGTCGACGCTCGAGATTTCCGGCGAGCCAACCACTGAGGATCTGTACTTTCAGAGCGATAACGCGATCGCCATGCCCCGGAGGGCGGAGAACTGGGACGAGGCCGAGGTAGGCGCGGAGGAGGCAGGCGTCGAAGAGTACGGCCCTGAAGAAGACGGCGGGGAGGAGTCGGGCGCCGAGGAGTCCGGCCCGGAAGAGTCCGGCCCGGAGGAACTGGGCGCCGAGGAGGAGATGGAGGCCGGGCGGCCGCGGCCCGTGCTGCGCTCGGTGAACTCGCGCGAGCCCTCCCAGGTCATCTTCTGCAATCGCAGTCCGCGCGTCGTGCTGCCCGTATGGCTCAACTTCGACGGCGAGCCGCAGCCCTACCCAACGCTGCCGCCTGGCACGGGCCGCCGCATCCACAGCTTCCGAGGTCACCTTTGGCTCTTCAGAGATGCAGGGACACACGATGGGCTTCTGGTTAACCAAACTGAATTATTTGTGCCATCTCTCAATGTTGACGGACAGCCTATTTTTGCCAATATCACACTGCCAGTGTATACTCTGAAAGAGCGATGCCTCCAGGTTGTCCGGAGCCTAGTCAAGCCTGAGAATTACAGGAGACTGGACATCGTCAGGTCGCTCTACGAAGATCTGGAAGACCACCCAAATGTGCAGAAAGACCTGGAGCGGCTGACACAGGAGCGCATTGCACATCAACGGATGGGAGATGTTTAA

nLuc-SMARCA2

ATGGTCTTCACACTCGAAGATTTCGTTGGGGACTGGCGACAGACAGCCGGCTACAACCTGGACCAAGTCCTTGAACAGGGAGGTGTGTCCAGTTTGTTTCAGAATCTCGGGGTGTCCGTAACTCCGATCCAAAGGATTGTCCTGAGCGGTGAAAATGGGCTGAAGATCGACATCCATGTCATCATCCCGTATGAAGGTCTGAGCGGCGACCAAATGGGCCAGATCGAAAAAATTTTTAAGGTGGTGTACCCTGTGGATGATCATCACTTTAAGGTGATCCTGCACTATGGCACACTGGTAATCGACGGGGTTACGCCGAACATGATCGACTATTTCGGACGGCCGTATGAAGGCATCGCCGTGTTCGACGGCAAAAAGATCACTGTAACAGGGACCCTGTGGAACGGCAACAAAATTATCGACGAGCGCCTGATCAACCCCGACGGCTCCCTGCTGTTCCGAGTAACCATCAACGGAGTGACCGGCTGGCGGCTGTGCGAACGCATTCTGGCGGGCTCGAGCGGCGCGATCGCCGCTGAGAAACTGTCACCAAATCCCCCCAAACTGACAAAGCAGATGAACGCTATCATCGATACTGTGATAAACTACAAAGATAGGTGTAACGTGGAGAAGGTGCCCAGTAATTCTCAGTTGGAAATAGAAGGAAACAGTTCAGGGCGACAGCTCAGTGAAGTCTTCATTCAGTTACCTTCAAGGAAAGAATTACCAGAATACTATGAATTAATTAGGAAGCCAGTGGATTTCAAAAAAATAAAGGAAAGGATTCGTAATCATAAGTACCGGAGCCTAGGCGACCTGGAGAAGGATGTCATGCTTCTCTGTCACAACGCTCAGACGTTCAACCTGGAGGGATCCCAGATCTATGAAGACTCCATCGTCTTACAGTCAGTGTTTAAGAGTGCCCGGCAGAAAATTGCCAAAGAGGAAGAGTAA

nLuc-SMARCA2 E1420S

ATGGTCTTCACACTCGAAGATTTCGTTGGGGACTGGCGACAGACAGCCGGCTACAACCTGGACCAAGTCCTTGAACAGGGAGGTGTGTCCAGTTTGTTTCAGAATCTCGGGGTGTCCGTAACTCCGATCCAAAGGATTGTCCTGAGCGGTGAAAATGGGCTGAAGATCGACATCCATGTCATCATCCCGTATGAAGGTCTGAGCGGCGACCAAATGGGCCAGATCGAAAAAATTTTTAAGGTGGTGTACCCTGTGGATGATCATCACTTTAAGGTGATCCTGCACTATGGCACACTGGTAATCGACGGGGTTACGCCGAACATGATCGACTATTTCGGACGGCCGTATGAAGGCATCGCCGTGTTCGACGGCAAAAAGATCACTGTAACAGGGACCCTGTGGAACGGCAACAAAATTATCGACGAGCGCCTGATCAACCCCGACGGCTCCCTGCTGTTCCGAGTAACCATCAACGGAGTGACCGGCTGGCGGCTGTGCGAACGCATTCTGGCGGGCTCGAGCGGCGCGATCGCCGCTGAGAAACTGTCACCAAATCCCCCCAAACTGACAAAGCAGATGAACGCTATCATCGATACTGTGATAAACTACAAAGATAGGTGTAACGTGGAGAAGGTGCCCAGTAATTCTCAGTTGGAAATAGAAGGAAACAGTTCAGGGCGACAGCTCAGTGAAGTCTTCATTCAGTTACCTTCAAGGAAAGAATTACCATCATACTATGAATTAATTAGGAAGCCAGTGGATTTCAAAAAAATAAAGGAAAGGATTCGTAATCATAAGTACCGGAGCCTAGGCGACCTGGAGAAGGATGTCATGCTTCTCTGTCACAACGCTCAGACGTTCAACCTGGAGGGATCCCAGATCTATGAAGACTCCATCGTCTTACAGTCAGTGTTTAAGAGTGCCCGGCAGAAAATTGCCAAAGAGGAAGAGTAA

nLuc-SMARCA2 L1465S

ATGGTCTTCACACTCGAAGATTTCGTTGGGGACTGGCGACAGACAGCCGGCTACAACCTGGACCAAGTCCTTGAACAGGGAGGTGTGTCCAGTTTGTTTCAGAATCTCGGGGTGTCCGTAACTCCGATCCAAAGGATTGTCCTGAGCGGTGAAAATGGGCTGAAGATCGACATCCATGTCATCATCCCGTATGAAGGTCTGAGCGGCGACCAAATGGGCCAGATCGAAAAAATTTTTAAGGTGGTGTACCCTGTGGATGATCATCACTTTAAGGTGATCCTGCACTATGGCACACTGGTAATCGACGGGGTTACGCCGAACATGATCGACTATTTCGGACGGCCGTATGAAGGCATCGCCGTGTTCGACGGCAAAAAGATCACTGTAACAGGGACCCTGTGGAACGGCAACAAAATTATCGACGAGCGCCTGATCAACCCCGACGGCTCCCTGCTGTTCCGAGTAACCATCAACGGAGTGACCGGCTGGCGGCTGTGCGAACGCATTCTGGCGGGCTCGAGCGGCGCGATCGCCGCTGAGAAACTGTCACCAAATCCCCCCAAACTGACAAAGCAGATGAACGCTATCATCGATACTGTGATAAACTACAAAGATAGGTGTAACGTGGAGAAGGTGCCCAGTAATTCTCAGTTGGAAATAGAAGGAAACAGTTCAGGGCGACAGCTCAGTGAAGTCTTCATTCAGTTACCTTCAAGGAAAGAATTACCAGAATACTATGAATTAATTAGGAAGCCAGTGGATTTCAAAAAAATAAAGGAAAGGATTCGTAATCATAAGTACCGGAGCCTAGGCGACCTGGAGAAGGATGTCATGCTTCTCTGTCACAACGCTCAGACGTTCAACTCGGAGGGATCCCAGATCTATGAAGACTCCATCGTCTTACAGTCAGTGTTTAAGAGTGCCCGGCAGAAAATTGCCAAAGAGGAAGAGTAA

EloB

ATGGACGTGTTCCTCATGATCCGGCGCCACAAGACCACCATCTTCACGGACGCCAAGGAGTCCAGCACGGTGTTCGAACTGAAGCGCATCGTCGAGGGCATCCTCAAGCGGCCTCCTGACGAGCAGCGGCTGTACAAGGATGACCAACTCTTGGATGATGGCAAGACACTGGGCGAGTGTGGCTTCACCAGTCAAACAGCACGGCCACAGGCCCCAGCCACAGTGGGGCTGGCCTTCCGGGCAGATGACACCTTTGAGGCCCTGTGCATCGAGCCGTTTTCCAGCCCGCCAGAGCTGCCCGATGTGATGAAGCCCCAGGACTCGGGAAGCAGTGCCAATGAACAAGCCGTGCACCTGCATGTCCACTCCCAGACGATGGCCAAGAGCAGAAACACAAGCTGGAGCCAGTGTCCTGGTTTGACAGCATGTTCAACGAGGGAACCCCAAGACGGACCCACACAGGTCCACCCACGCTGGGGGCTGTAA

EloC

ATGGATGGAGAGGAGAAAACCTATGGTGGCTGTGAAGGACCTGATGCCATGTATGTCAAATTGATATCATCTGATGGCCATGAATTTATTGTAAAAAGAGAACATGCATTAACATCAGGCACGATAAAAGCCATGTTGAGTGGCCCAGGTCAGTTTGCTGAGAACGAAACCAATGAGGTCAATTTTAGAGAGATACCTTCACATGTGCTATCGAAAGTATGCATGTATTTTACGTACAAGGTTCGCTACACTAACAGCTCCACCGAGATTCCTGAATTCCCAATTGCACCTGAAATTGCACTGGAACTGCTGATGGCTGCGAACTTCTTAGATTGTTAA
